# Deep learning assessment of nativeness and pairing likelihood for antibody and nanobody design with AbNatiV2

**DOI:** 10.1101/2025.10.31.685806

**Authors:** Aubin Ramon, Niccolò Frassetto, Haowen Zhao, Xing Xu, Matthew Greenig, Shimobi Onuoha, Pietro Sormanni

## Abstract

Immune systems create antibodies that balance good binding and stability with low toxicity and self-reactivity. Quantifying the nativeness of a candidate sequence – its likelihood of belonging to natural immune repertoires – has thus emerged as a valuable strategy for hit selection from synthetic libraries, optimisation and humanisation, and for guiding de novo design towards developable candidates. We previously introduced AbNatiV, a transformer-based VQ-VAE for nativeness assessment, which proved effective across multiple nanobody engineering tasks. However, AbNatiV1 operated on unpaired sequences, limiting applicability to conventional VH-VL antibodies. Moreover, its performance on nanobody nativeness was constrained by the limited number and diversity of nanobody repertoires available at the time. Here, we sequenced new camelid repertoires, curated additional recent datasets, and present AbNatiV2: an enhanced architecture comprising various models each trained on ≥ 20 million sequences. AbNatiV2 improves nanobody nativeness classification across held-out and diverse test sets, and more robustly detects nativeness changes upon CDR grafting. We also introduce p-AbNatiV2, a cross-attention model fine-tuned on 3.7 million paired human sequences. p-AbNatiV2 provides residue- and sequence-level humanness for VH/VL pairs and learns pairing-likelihood via noise-contrastive training. On held-out tests, it assigns the native pair a higher score in 74% of cases, substantially outperforming recent pairing models. Together, AbNatiV2 and p-AbNatiV2 extend nativeness assessment and engineering to both nanobodies and conventional antibodies, supporting design decisions at single-residue, Fv-sequence, and paired-domain levels. We make AbNatiV2 available as downloadable software and webserver.

## 2 Introduction

The immune system operates as a highly optimised antibody-discovery platform, typically producing antibodies combining good binding and stability with low toxicity, long half-life, and no cross-reactivity with self-antigens. Antibodies designed to resemble those naturally generated by the immune system are therefore more likely to retain these favourable in vivo properties, which are incredibly hard to measure experimentally in good throughput and therefore to directly predict computationally. Quantifying the likelihood that a given sequence belongs to the distribution of immune-system-derived antibodies (i.e., its nativeness) is thus an effective strategy for designing or refining candidate antibody therapeutics. This principle has long underpinned the humanisation of murine antibodies to reduce immunogenicity by maximising their humanness (1). More recently, nativeness assessment has been extended to camelid nanobodies (V_H_Hs), where evaluating the V_H_H-nativeness facilitates the selection and engineering of nanobodies whose stability, solubility, and structural features reflect those found in camelid nanobody repertoires (2). Given the expanding use of nanobodies across a range of therapeutic applications (3) (e.g., to create multi-specific formats (4, 5), as antigen receptors in CAR-T cell therapies (6, 7) and for nanoparticle targeting (8–10)), the demand for reliable nanobody engineering continues to rise.

Beyond hit selection and optimisation, the computational assessment of nativeness offers unique value for antibody de novo design (11). While in silico generation enables an extensive exploration of sequence space, it can produce sequences that deviate from evolutionary distributions and that often show poor biophysical viability in vitro and likely poor in vivo properties. For instance, the generative model RFdiffusion (12) was successfully used to generate libraries of single-chain antibodies (scFvs), which however showed a rather low surface expression yield despite the use of fixed framework regions (13). Nativeness assessment can help filter out unrealistic candidates and guide the generation of more developable antibody sequences (14–16).

Fuelled by the steady expansion of antibody repertoire data from next-generation sequencing (NGS), antibody-specific protein language models (PLMs) have gained increasing popularity. The Observed Antibody Space (OAS) database (17) currently contains around two billion sequences. Building on such large-scale resources, antibody-specific PLMs like AntiBERty (18), AbLang2 (19), p-IgGen (20) and Humatch (21) enable sequence generation for library construction, offer rich embeddings to streamline downstream predictions (22), and provide sequence pseudo-loglikelihood scores that can be leveraged for zero-shot predictions and antibody selection or optimisation (23–25). Dedicated nanobody PLMs such as nanoBERT (26) have also been developed, although nanobody sequence datasets remain comparatively limited.

Whereas PLMs excel at sequence generation and provide evolutionarily informed embeddings, the AbNatiV model (2) was introduced with a different emphasis: the direct and interpretable assessment of nativeness at both full sequence and single residue resolution. AbNatiV is a transformer-based vector-quantised variational auto-encoder (VQ-VAE) model designed to quantify how far any given sequence lies from the learned distribution of native antibody sequences. Unlike PLMS that rely on cross-entropy-based likelihoods, AbNatiV employs masked learning and a mean squared error loss, yielding a direct distance measure from the learnt native space. Besides, by mapping sequences into a limited set of discrete latent codes in an information bottleneck, AbNatiV’s VQ-VAE framework effectively captures recurring motifs and long-range interactions (2).

Since its publication, AbNatiV has been adopted by multiple research groups (14, 27–29). It was notably employed to humanise a nanobody used as the chimeric antigen receptor (CAR) in a CAR T-cell construct, which demonstrated superior ex vivo selectivity over its parental nanobody as well as preclinical efficacy in vivo in murine models of glioblastoma (29). Additionally, AbNatiV has been integrated with other predictors to simultaneously optimise multiple properties (including solubility, thermostability, humanness, and V_H_H-nativeness), ultimately enabling the engineering of multiple improved nanobodies functional in the brain (27, 28). Beyond hit optimisation, AbNatiV was also used to guide a diffusion model for generating antibody scaffolds with improved humanness and nativeness, while preserving binding affinity and functionality in vitro (14).

Notwithstanding these advances, AbNatiV1 showed limitations in nanobody nativeness assessment. While the human models perfectly distinguished human antibody sequences from artificial PSSM-derived sequences that are devoid of long range interactions, performance was lower for the nanobody model (2), pointing to limitations in the nanobody training corpus and motivating the expansion carried out in this study.

Moreover, the human models were trained only on unpaired heavy and light chains. Although large-scale paired sequencing still relies on laborious single-cell approaches (30), new resources such as PairedAbNGS (31) and expanded OAS datasets are beginning to bridge this gap. Recent models, including p-IgGen (20), Humatch (21) and ImmunoMatch (32), have begun to leverage paired sequences in their training. Incorporating paired data into nativeness assessment is important, as correct VH–VL pairing underpins the structural integrity and stability of Fv regions (33). Failure to account for pairing can compromise synthetic library quality (34) and reduce the functionality of bispecific formats, especially in common light chain platforms (35, 36).

In this work, we expanded the nanobody training corpus to 21 million V_H_H sequences by curating recently published repertoires (37, 38) and adding around 9.2 million nanobodies sequenced in-house. The model’s architecture is scaled up in size and upgraded with modern transformer components such as rotary positional encodings, SwiGLU transition layers, and attention gating mechanisms. A focal reconstruction loss is also introduced to mitigate germline biases. We show that AbNatiV2 consistently improved V_H_H nativeness predictions, generalised to newly sequenced repertoires where AbNatiV1 underperforms, and more faithfully detected changes in nativeness after CDR grafting. For human antibodies, we scale the models and train them on a tenfold larger dataset. A unified VL model spanning Vκ and Vλ improved discrimination against other species, with the largest gains observed against rhesus light chains.

Building on these unpaired models, we introduce p-AbNatiV2, a cross-attention VQ-VAE fine-tuned on >3.7 million unique paired human sequences. It (i) scores humanness jointly for VH and VL at paired-domain, individual sequence, and single-residue resolution, and (ii) predicts pairing likelihood via noise-contrastive learning. This enables to perform tasks that AbNatiV1 could not accomplish: ranking light-chain partners for a given heavy chain, triaging designs that disrupt pairing, and prioritising candidates and engineering mutations in common light-chain formats or other pairing tasks. On held-out tests, p-AbNatiV2 assigned higher likelihood to the native partner in 74% of one-vs-one comparisons and ranked the native pair in the top five in >35% of cases in one-vs-fifty comparisons, substantially outperforming the current state of the art. We release AbNatiV2 and p-AbNatiV2 as source code and via a user-friendly web server, together with the underlying sequence datasets, to support paired-antibody design and evaluation.

## 3 Results

### 3.1 The unpaired AbNatiV2 models

AbNatiV2 is a transformer-based vector-quantised variational auto-encoder (VQ-VAE) (39) trained via self-supervised masked learning on aligned antibody sequences (see **Fig. 1a**). The encoder compresses one-hot encoded amino acid sequences into a latent transformer block using a 1D convolutional patch. Within the transformer layers, a rotary positional embedding encodes for relative positional information prior to the attention mechanism (40). The output of the attention dot-product is dynamically modulated by an integrated gated mechanism (41), and processed through a residual connection with a SwiGLU activation mechanism (42). Finally, the encoder’s output is projected onto a discrete codebook via a nearest neighbour lookup and reconstructed by a decoder that mirrors the encoder. At inference time, the nativeness of an antibody sequence is derived from the reconstruction mean-squared error (MSE) calculated between the original input and the reconstructed output (see Methods).

**Fig. 1.**
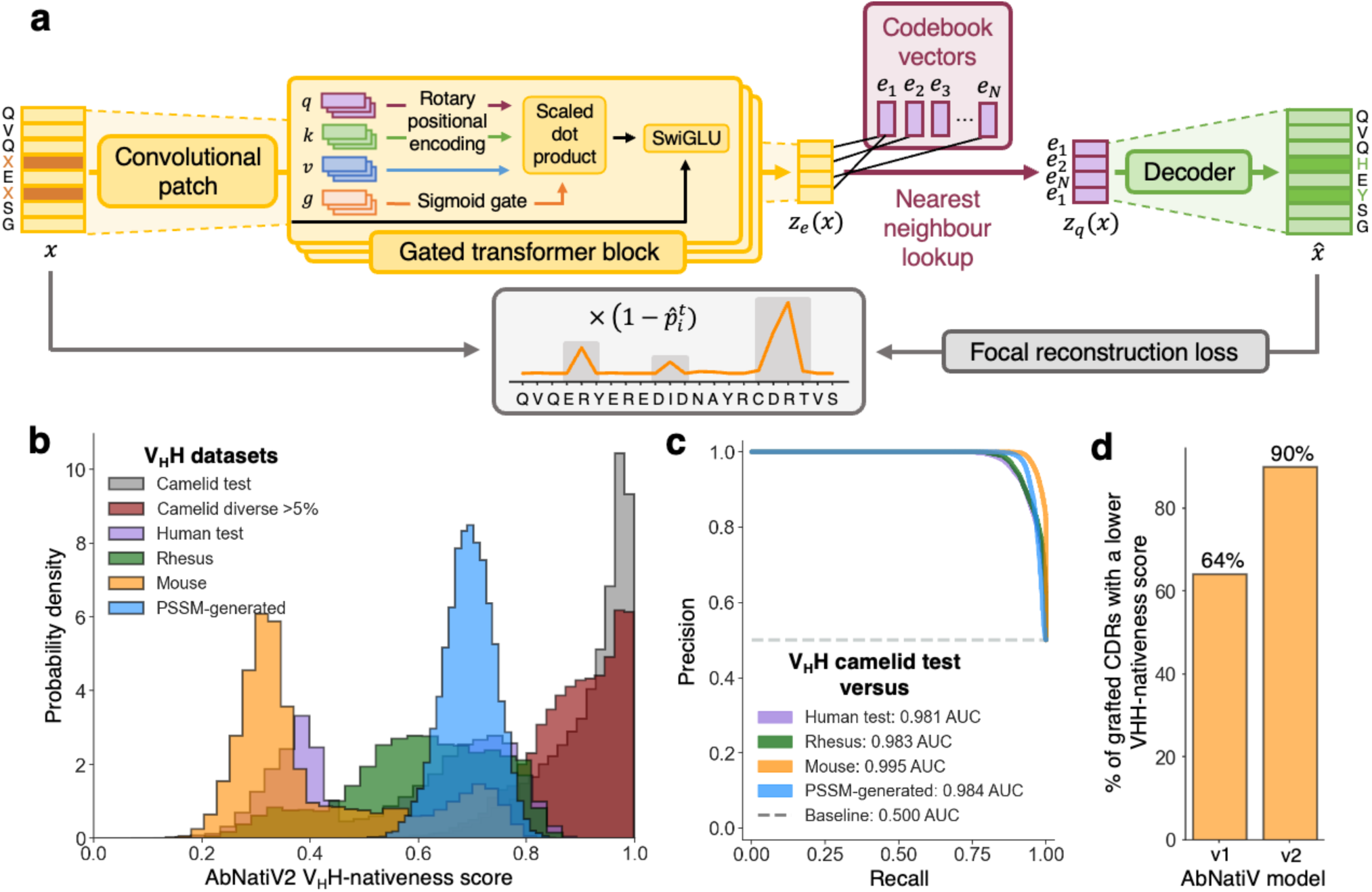
The unpaired AbNatiV2 model and nanobody nativeness predictions. (**a**) Architecture of the AbNatiV2 model. Through the encoder (in yellow), the input sequence 𝑥 is compressed into a latent transformer block via a convolutional patch. In the latent space (in burgundy), 𝑧_𝑒_(𝑥) is projected into the discrete embedding 𝑧_𝑞_(𝑥) with a nearest neighbour lookup on a codebook 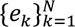 of 𝑁 code vectors. Finally, the output *x̂* is reconstructed by a decoder (in green), mirroring the encoder architecture. During training, a portion of input residues is replaced with masked vectors (in a darker shade). The training loss combines a vectorisation commitment term (see Methods) with a focal reconstruction loss (FRL; in grey) which modulates the positional mean-squared error (MSE) of each residue position 𝑖 by 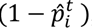, where 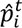 represents the output likelihood assigned to the target (ground-truth) residue in *x̂* at position 𝑖. (**b**) The AbNatiV2 V_H_H-nativeness score distributions of various test sets (see legend); the V_H_H camelid test is a random split of unique sequences, while the camelid diverse has sequences at least 5% different from the closest training set sequence. The V_H_H PSSM-generated database is made of artificial sequences randomly generated using residue positional frequencies from the PSSM of the V_H_H sequences (**Supplementary Fig. 3a**). Each dataset contains 10,000 sequences. (**c**) Plots of the PR curves used to quantify the ability of AbNatiV2 to distinguish the V_H_H camelid test set from the other datasets (see legend, which also reports the AUC values). The baseline (dashed line) corresponds to the performance of a random classifier. The corresponding plot against the V_H_H diverse set is in **Supplementary Figure 7a**. (**d**) All three CDRs from test set sequences are computationally grafted onto the universal V_H_H framework (UF; see Methods). The bar plot shows that 90% of them have a lower CDR V_H_H-nativeness score when grafted onto the UF than when they are within their native framework, compared to 64% for AbNatiV1.

First, we assessed whether the changes we made to the architecture and to the loss function translate into an improved performance over AbNatiV1. To this end, we worked using the same training and validation datasets, as well as the same hyper-parameters and singly introduced architectural modifications and different loss functions (**Supplementary Fig. 1**). We find that the use of rotary embeddings for encoding relative positional information, combined with the enhanced representation capacity of gated SwiGLU activation functions, resulted in a significant performance improvement, with a nearly 5% gain in reconstruction loss. These results confirmed the advantage of these architectural improvements (40, 42, 43) for sequence-based protein models.

Two alternative reconstruction loss strategies were explored to address the limitations of AbNatiV1 in capturing some high order relationships between positions and effectively distinguishing native V_H_H sequences from artificial sequences (see Methods and **Supplementary Figure 2**). Such artificial sequences are generated using the positional residue frequency derived from the training set sequences (**Supplementary Fig. 3a**), but with residue positions filled independently of each other and thus devoid of higher-order relationships beyond random expectations. Building upon the focal loss approach described in Ref. (44) and recently employed by AbLang2 to tackle germline bias in PLMs (19), the standard mean-squared error (MSE) used as reconstruction loss in AbNatiV1 is replaced with a focal reconstruction loss (FRL) still based on MSE, but that dynamically down-weights positions the model reconstructs with high accuracy (see **Figure 1a** and Methods for more details). By mitigating the loss contribution of these easy-to-predict positions, the FRL drives the model to allocate more learning capacity to hard-to-predict residues, which may be part of hypervariable regions (e.g., H-CDR3) or the result of somatic mutations. To enable the prediction of these positions, this approach encourages the model to capture high order relationships between residues or sequence regions inherent to native antibodies. In a preliminary training on camelid sequences, the FRL effectively improved the model’s ability to distinguish native camelid sequences from PSSM-generated sequences (PR-AUC = 0.954 for FRL vs. 0.945 for MSE after 24 epochs, while also improving the reconstruction accuracy by ∼1% (**Supplementary Fig. 2a-b**).

In parallel, a static version of the FRL (sFRL) was tested by replacing the dynamic focal down-weighting with a fixed conservation index penalty, reflecting the conservation of each sequence position in the AHo alignment (see Methods). The sFRL aims to specifically direct the model’s focus on more variable regions. While this approach further improved the model’s ability to discriminate native sequences from PSSM-generated ones (PR-AUC = 0.965), its static nature resulted in nearly 2% decline in reconstruction accuracy (**Supplementary Fig. 2**). This trade-off resulted in a drop of discrimination accuracy between V_H_Hs and VHs of other species, such as mouse (PR-AUC = 0.955 for sFRR vs. 0.986 for FRL; **Supplementary Figure 2c**). Consequently, the dynamic FRL was adopted as the reconstruction loss for AbNatiV2.

### 3.2 Nanobody nativeness assessment

As a rapidly expanding class of biologics, nanobodies are currently benefitting from a substantial surge in experimental data released in the literature (3). Over the span of two years, we significantly expanded the original AbNatiV1 training dataset, which contained ∼2 million sequences from very few sources, with 18 million additional sequences derived from newly published camelid repertoires (37, 38), and novel non-immune alpaca libraries sequenced for this study (**Supplementary Table 1**). The updated dataset comprises V_H_H sequences derived from alpaca (77%), llama (16%), and camel (7%) repertoires (**Supplementary Fig. 4**). All sequences were aligned on the AHo numbering scheme (45) (see Methods).

The AbNatiV2 model, trained on this expanded dataset of 20,208,125 nanobody sequences, employs self-supervised masked learning to map antibody sequences into a latent space tuned on immune-derived nanobodies. This space captures evolutionary and biophysical features intrinsic to native nanobodies, such as their ability to fold independently of a VL counterpart. The model ultimately provides a V_H_H-nativeness score, quantifying the resemblance of any antibody sequence to native single-domain antibodies from camelid repertoires. Nativeness scores are linearly rescaled to approach 1 for highly native sequences, with 0.8 serving as the optimal threshold to distinguish native nanobody sequences from non-native ones (see Methods).

The hyperparameter search resulted in an upscaling of the model compared to AbNatiV1, tailored to effectively harness the expanded training dataset. The number of transformer layers and their embedding size are increased, yielding approximatively 92 million free parameters, representing a substantial expansion from the 17 million parameters in AbNatiV1 (**Supplementary Table 2**). The model was trained for 30 epochs, with each epoch taking approximately 5 hours to complete on a single GPU (NVIDIA A100-SXM-80GB). Training was terminated upon convergence (**Supplementary Fig. 5a**), while controlling for overfitting on the validation set (**Supplementary Fig. 6a**).

By integrating the extended nanobody training dataset and updated architectural features, AbNatiV2 achieved notable performance improvements over version 1 (**Fig. 1b-d, Supplementary Fig. 7a, Table 1**). In the V_H_H-AbNatiV2 model, PSSM-generated sequences were substantially shifted toward lower nativeness scores (**Fig. 1b** and **c**, PR-AUC = 0.984 for AbNatiV2 vs 0.961 for AbNatiV1). In parallel, the model maintained high PR-AUC classification accuracy against human, rhesus, and mouse datasets (**Fig. 1a-b**), while increasing sequence reconstruction accuracy (**Table 1**). Furthermore, AbNatiV2 retained classification performance on a diverse test set of sequences, selected to be at least 5% distant from any training sequence (see Methods). When distinguishing these diverse test sequences from artificial PSSM-generated ones, AbNatiV2 showed a PR-AUC of 0.977 while AbNatiV1 drops to 0.926 (**Table 1** and **Supplementary Fig. 7a**).

**Table 1.** Performance comparison between the unpaired AbNatiV1 and AbNatiV2 models. . The assessment was carried out for AbNatiV2 (shaded rows) and AbNatiV1 (unshaded rows), trained on camelid V_H_H (rows 1- 2), human VH (rows 3-4) and human VL (rows 5-7) antibody sequences. Central columns display the area under the PR curve (PR-AUC; shown in **Fig. 1c** and **Supplementary Fig. 7 and 9**) evaluating the ability of the models to distinguish their respective test (T) and diverse test (D) set sequences from human, rhesus, mouse and PSSM-generated sequences (column headers). For the AbNatiV1 models, their original test and diverse sets were used to compute the performances on the new human, rhesus and mouse sets. The last two columns report each model’s ability to reconstruct their respective test (T) and diverse test (D) set sequences. For the light chains, AbNatiV1 models were separately trained on Vκ and Vλ sequences and the performances are reported on each Vκ or Vλ subpopulation of each benchmarking set.

| Chain type | AbNatiV Model | Classification (PR-AUC) ↑<br>against |  |  |  |  |  |  |  | Re-<br>construction<br>accuracy<br>(%) ↑ |  |
| --- | --- | --- | --- | --- | --- | --- | --- | --- | --- | --- | --- |
|  |  | Human |  | Rhesus |  | Mouse |  | PSSM-generated |  | T | D |
|  |  | T | D | T | D | T | D | T | D |  |  |
| Camelid<br>V<sub>H</sub>H | V2 | 0.981 | 0.972 | 0.983 | 0.975 | 0.995 | 0.993 | 0.984 | 0.977 | 96.6 | 95.5 |
|  | V1 | 0.965 | 0.922 | 0.976 | 0.945 | 0.994 | 0.985 | 0.961 | 0.926 | 95.4 | 95.4 |
| Human<br>VH | V2 | N/A | N/A | 0.965 | 0.937 | 0.999 | 0.997 | 1.000 | 1.000 | 97.4 | 96.5 |
|  | V1 | N/A | N/A | 0.950 | 0.893 | 0.994 | 0.983 | 0.999 | 0.996 | 96.0 | 93.5 |
| Human<br>VL | V2 | N/A | N/A | 0.949 | 0.851 | 0.998 | 0.992 | 0.996 | 0.987 | 98.7 | 97.8 |
|  | V<sub>κ</sub>-V1 | N/A | N/A | 0.808 | 0.768 | 0.998 | 0.996 | 0.977 | 0.967 | 98.2 | 97.9 |
|  | V<sub>λ</sub>-V1 | N/A | N/A | 0.851 | 0.802 | 1.000 | 1.000 | 0.982 | 0.968 | 98.3 | 97.8 |

Although the original AbNatiV1 maintained strong performances on its own test and diverse sets (**Table 1**), it struggled on the new V_H_H datasets that contain many more sequences: more than half of these sequences are scored below 0.8 (**Supplementary Fig. 8a**). Using MMseqs clustering with a 90% sequence identity cutoff, the AbNatiV1 sequences covered only around 1.2M of the 12M clusters present in the sequence corpus of AbNatiV2. These findings indicate that the AbNatiV1’s nanobody nativeness assessment was constrained by the limited size and diversity of its original training dataset, resulting in poor generalisability to new sequenced repertoires (2). In contrast, AbNatiV2 offers a more robust nativeness assessment built on a substantially bigger and much more diverse nanobody dataset.

AbNatiV2 was also able to detect nativeness loss in CDR loops grafted onto a different scaffold, offering valuable insights for CDR-grafting strategies – a widely used approach to engineer nanobodies with reduced immunogenicity or enhanced developability (46–50). The model consistently assigned lower nativeness scores to the CDRs of all six nanobodies that exhibited binding loss upon grafting onto the universal scaffold (UF) in Ref. (47) (**Supplementary Fig. 7b**). Additionally, AbNatiV2 predicted lower nativeness for 90% of CDRs grafted onto the same UF scaffold across 10,000 test sequences, outperforming AbNatiV1, which detected a nativeness loss in 64% of cases on the same test sequences (**Fig. 1d**), compared to 86% on its own test set (2).

Taken together, these findings highlight how the integration of an expanded training dataset, tailored learning strategies, and architectural enhancements can substantially improve the nativeness assessment of nanobody sequences. The biggest improvement was observed in the model’s ability to capture high-order positional relationships, enabling AbNatiV2 to distinguish more effectively native nanobody sequences from artificial ones, and to determine more accurately if CDR loops are grafted in the right context. Furthermore, it significantly increased the model’s capacity to generalise to diverse sequences that are distant from the training set.

### 3.3 Humanness assessment of unpaired antibody sequences

AbNatiV2 was separately trained on unpaired heavy-chain and light-chain sequences derived from human immune systems. Previously, AbNatiV1 was trained on a random subset of 2 million sequences from the OAS. Following the upscaling efforts undertaken for the nanobody model, we performed a more rigorous parsing of the entire OAS dataset, ultimately curating a highly diverse set of 19,654,973 VH and 21,228,567 VL sequences (see Methods). The VH and VL models were trained for 35 epochs with the same hyperparameters used for the V_H_H model. Training was terminated upon convergence (**Supplementary Fig. 5b-c**), while controlling for overfitting (**Supplementary Fig. 6b-c**).

These models quantified the nativeness of human antibody sequences, referred to as ‘humanness’ (**Fig. 2a–b**). Although human-VH-AbNatiV1 already performed very well, VH-AbNatiV2 further increased PR–AUC for most human-versus-other-datasets comparisons (**Table 1** and **Supplementary Fig. 9**). Performance gains were larger when using the human diverse test set, suggesting better generalisation to sequences more distant from the training distribution (**Table 1**). For example, in discriminating human from rhesus VH sequences, AbNatiV1 dropped from a PR-AUC of 0.950 to 0.893 when moving from the human to the human-diverse test set, whereas AbNatiV2 changed from 0.965 to 0.937, denoting both higher performance and smaller degradation.

**Fig. 2.**
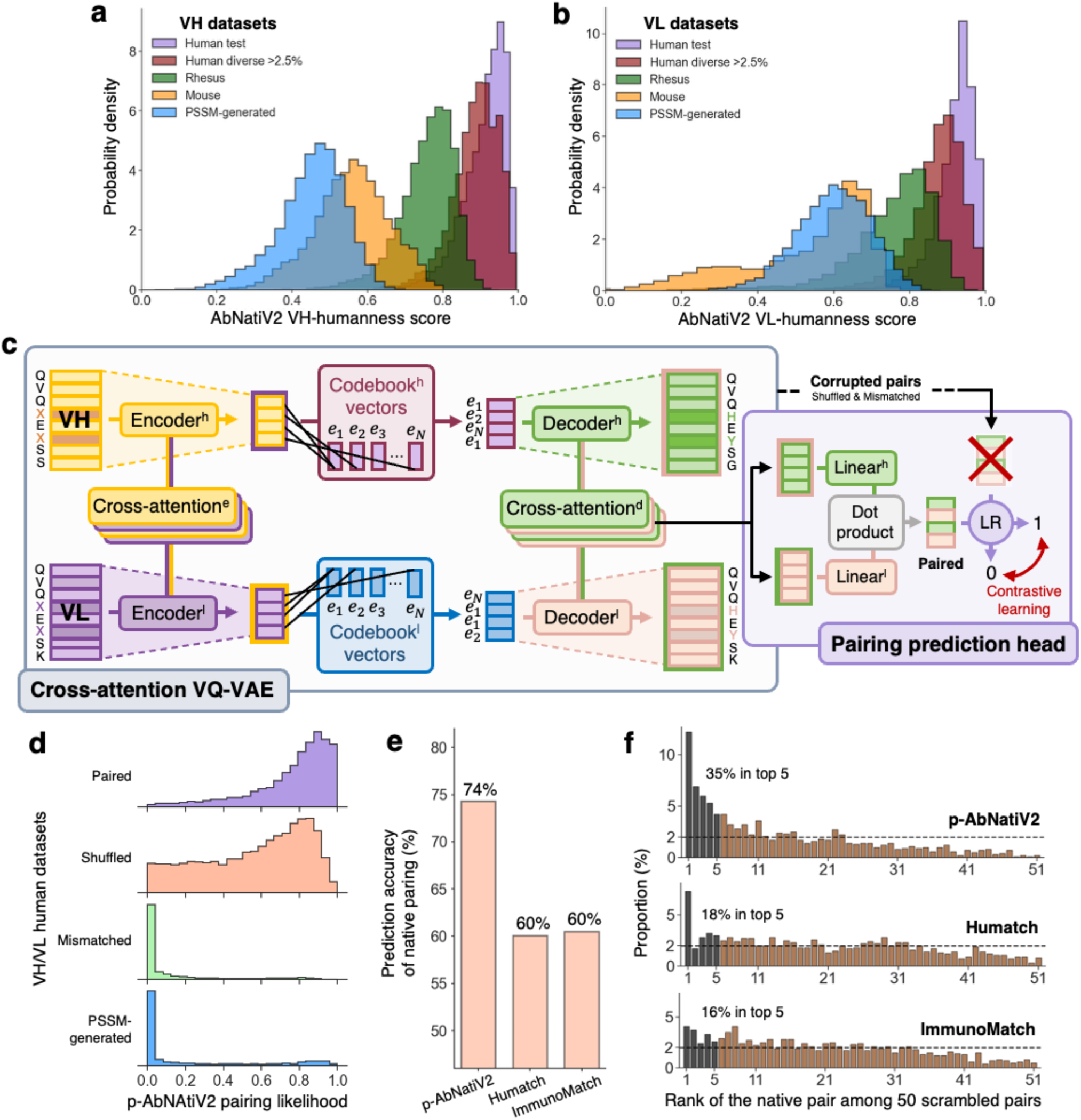
The paired p-AbNatiV2 model and antibody humanness. The AbNatiV2 human VH (**a**) and VL (**b**) humanness score distributions of the human test (in purple), human diverse (in red), rhesus (in green), mouse (in orange), and PSSM-generated (in blue) datasets. Each dataset contains 10,000 sequences. (**c**) The paired p- AbNatiV2 architecture is composed of two units: a cross-attention VQ-VAE (in the light grey box), which projects and reconstructs VH (top row) and VL (bottom row) sequences conjointly via cross-attention layers, and a pairing prediction head (in the light pink box), which classifies natively paired sequences from unpaired ones via noise-contrastive learning (Methods). (**d**) Distributions of p-AbNatiV2 pairing likelihood scores for 10,000 native human test pairs (in purple), shuffled pairs generated by scrambling the light chains across the different native pairs (in salmon), mismatched human sequences generated by mixing heavy and light chains from different B-cell classes (in green) and studies, and artificial PSSM-generated sequences (in blue, see Methods; corresponding distributions for Humatch and ImmunoMatch are in **Supplementary Figure 14)**. (**e**) Pairing prediction accuracy of p-AbNatiV2 (74.3%) compared with Humatch (60.1%) and ImmunoMatch (60.5%), assessing the percentage of times in which the pairing likelihood of a human VH paired with its native VL is higher than that of the same VH paired with a random VL within the same somatic maturation bin. This evaluation included 10,000 VHs from the VH/VL paired test set. (**f**) Ranking distributions of the native human VH-VL pair among 50 shuffled pairs within the same somatic maturation bin (for N=1,000 different pairs) for p-AbNatiV2 (top), Humatch (middle) and ImmunoMatch (bottom). Black bars highlight the proportion of instances in which the native pair ranked within the top 5 for each method (35% for p-AbNatiV2, 18% for Humatch and 16% for ImmunoMatch). The horizontal line at 2% indicates the expected uniform distribution of a random classifier.

While for AbNatiV1 we introduced separate Vκ and Vλ models, for AbNatiV2 we develop a single VL model trained on a dataset that contains both Vκ and Vλ light chains. The reason is that this model also serves as a foundation for our paired VH/VL model. VL-AbNatiV2 demonstrated a considerable increase in classification accuracy against VL rhesus sequences (PR-AUC = 0.949 for VL-AbNatiV2 compared to 0.808 and 0.851 for Vκ- and Vλ-AbNatiV1, respectively) and small changes against other datasets (**Table 1**).

In conclusion, scaling up the VH and VL models further improved the accuracy and robustness of antibody humanness assessment. In particular, the unified light-chain model, together with the VH model, provide the pre-trained foundations for developing a VH/VL paired model combining the insights captured by each model individually with partner-aware humanness and pairing-likelihood scores.

### 3.4 The paired p-AbNatiV2 model

Assessing the humanness of unpaired antibody sequences is a powerful approach to screen for hits with high humanness in antibody discovery. However, engineering a lead antibody or designing one de novo without accounting for interactions between heavy and light chains risks introducing mutations that compromise functionality and developability, by weakening or disrupting chain pairing or inter- domain CDR interactions that stabilise paratopes (33–36). To address this challenge, we develop p- AbNatiV2 (**Fig. 2c**), a paired VQ-VAE that leverages the AbNatiV2 unpaired models to jointly evaluate the humanness of VH/VL pairs and predict their pairing compatibility.

Designed as a multimodal network (51), p-AbNatiV2 is built by combining the pre-trained unpaired models with cross-attention layers (52) in the encoder and decoder, interconnecting the VQ-VAE architectures (see Methods). To prevent overfitting of the considerably smaller and less diverse paired training set, the pre-trained weights of the unpaired models are kept constant and only the weights of the cross-attention layers are updated during training. A cross-gating mechanism restricts information flow between chains early in training to facilitate a smooth integration of the cross-attention layers. This architecture enables the joint processing of heavy and light chain sequences together while retaining learnings from the unpaired training.

On top of the cross-attention VQ-VAE, we integrate a pairing prediction head that predicts the pairing likelihood between the two input chains (i.e., the likelihood that the heavy and light chains form a native pair). Specifically, this prediction head combines the heavy and light outputs from the final cross-attention layer into a logistic regression (LR), trained via noise-contrastive learning with an additional loss function term. During the training of this prediction head, native sequence combinations are labelled as ‘paired,’ while two types of ‘mis-paired’ negatives are introduced: shuffled pairs generated by scrambling the light chains within the current training batch, and mismatched pairs generated by mixing heavy and light chains from different human B-cell classes, species, and studies. The contrastive pairing prediction loss is summed with the losses from each VQ-VAE, enabling the whole model to jointly learn the underlying distribution of paired sequences (see Methods). p- AbNatiV2 is designed to provide a humanness score for the VH/VL pair, two separate individual scores for VH and VL sequences, a residue-resolution humanness profile, and a VH/VL pairing likelihood score.

p-AbNatiV2 is fine-tuned on a novel dataset of 3,745,614 unique paired human sequences, constructed by combining the training dataset of p-IgGen (20), derived from the OAS (17), with the PairedAbNGS dataset (31) and filtering for unique paired sequences. All heavy and light chain sequences were aligned and processed identically to the unpaired models (Methods). Hyperparameters were manually tuned, and the pairing prediction loss was carefully scaled to prevent it from dominating the VQ-VAE reconstruction objective. To enable rapid adaptation of the newly initialised pairing head on the pretrained backbone, its learning rate was set to ten times that of the main VQ-VAE trunk. (**Supplementary Table 2**). Training was halted at 20 epochs (**Supplementary Fig. 5d**).

The humanness assessment from the main VQ-VAE trunk was tested on paired rat and mouse sequences from the OAS database alongside human test, diverse test, and PSSM-generated sequences (**Supplementary Fig. 10, Supplementary Table 3**). p-AbNatiV2 demonstrated nearly perfect classification performance, with PR-AUC values approaching 1 when distinguishing human paired sequences from non-human ones. When evaluating on the heavy and light chain sequences separately, p-AbNatiV2 exhibited slightly lower performance on VH sequences than VL sequences. It achieved a PR-AUC of 0.985 for VH rat and human diverse test sequences while reaching 0.999 for the corresponding VL sequences (see **Supplementary Table 3**). Additionally, p-AbNatiV2 achieved a high reconstruction accuracy of 96.9% on the test set, with a less than 1% decrease in accuracy on the diverse test set, underscoring the model’s ability to generalise despite some minor indications of overfitting during training (**Supplementary Figure 6d**). Finally, p-AbNatiV2 consistently outperformed its unpaired VH- and VL-only counterparts across all benchmarks, and notably on the diverse sets (see **Supplementary Table 3**).

We applied AbNatiV2 and p-AbNatiV2 humanness assessments to evaluate their relationship with immunogenicity in therapeutic antibodies. First, we assessed the ability of these models to classify 190 human-derived therapeutics from 342 non-human-origin therapeutics (mouse, chimeric, and humanized). For the unpaired AbNatiV2 models, the heavy and light chain humanness scores were averaged into a single score. For the paired p-AbNatiV2 model, the humanness score of the pair was used. We compared their performance with p-IgGen (20) and Humatch (21), two recent humanness assessment tools. Using methodologies from their respective publications, p-IgGen scores were computed as the perplexity of concatenated heavy and light chains, while Humatch scores were derived from the minimum score among heavy, light, and paired assessments. Resulting PR-AUC over 1,000 bootstrapping resamples are reported in **Supplementary Figure 11a-b**. p-IgGen (PR-AUC = 0.952 and Humatch (PR-AUC = 0.850) significantly underperformed compared to all AbNatiV models. The original AbNatiV1 model achieved the highest mean PR-AUC (0.976 ± 0.006 SE); however, the differences from the AbNatiV2 models and the OASis 9-mer peptide search (53) are within the standard errors from bootstrap, indicating no significant difference among these top performers.

Furthermore, correlation between the humanness assessments and clinical ADA immunogenicity values of 216 therapeutic antibodies were computed (**Supplementary Fig. 11b-c**). The Pearson’s correlations of both the unpaired (r= −0.47) and paired (r = −0.51) AbNAtiV2 models aligned to the best reported performances on this dataset (i.e., p-IgGen with r = 0.53 (20)). We note, however, that this is a highly heterogeneous and inconsistent dataset that combines datapoints from different clinical studies that were not designed to be comparable with each other (2). For instance, they differ remarkably in patient population, antibody dosage, and treatment length, among many other factors, suggesting that strong correlations between humanness and these data should not be expected.

Collectively, these findings demonstrate that the development of the paired p-AbNatiV2 model enables the robust humanness assessment of paired sequences.

### 3.5 Prediction of variable-domain pairing likelihood

In addition to the cross-attention VQ-VAE model that jointly evaluates the humanness of paired antibody sequences, p-AbNatiV2 incorporates a prediction head that estimates the pairing likelihood between VH and VL sequences. After 20 epochs, no overfitting was observed for the pairing likelihood assessment (**Supplementary Fig. 6e**).

The pairing prediction head is trained via noise-contrastive learning strategy (54), where the model learns to distinguish native VH-VL pairs (positives) from corrupted mis-paired examples (negatives). We generate two negative sets: (i) shuffled negatives by in-batch random reassignment of VLs, and (ii) mismatched negatives by pairing VH and VL chains from different human B-cell classes, species and studies, following an approach similar to Humatch’s (21). Mismatched pairs are typically far apart in evolutionary space and thus constitute ‘easy’ negatives for the model to spot. By contrast, shuffled pairs are likely to be much closer, making them ‘hard’ negatives. However, because a given VH can productively pair with multiple VLs and vice versa (34), shuffled pairs are also noisy negatives, as chances are that a good fraction of them would constitute perfectly productive pairs. To account for this asymmetry, we apply a weighted loss, penalising errors on mismatched and shuffled negatives with varying ratios (see **Supplementary Figure 12**). Ultimately, we found that using equal weights effectively limited the impact of plausible shuffled pairs (**Supplementary Fig. 12a**) while preserving strong separation from high-confidence mismatches (**Supplementary Fig. 12b**).

The resulting p-AbNatiV2 model clearly distinguished native test pairs from both mismatched pairs and PSSM-generated artificial sequences (**Fig. 2d**). As expected, given the noisy nature of shuffled negatives, pairing likelihood distributions between native test pairs and randomly shuffled ones exhibited greater overlap. The same behaviour was observed on the diverse paired test set, ruling out overfitting (**Supplementary Fig. 13**). To quantitatively assess the performances of this prediction head, we evaluate the model accuracy at distinguishing native pairs from randomly shuffled counterparts (**Fig. 2e-f**). For each VH, new VLs were randomly selected among those in the test set that exhibited a similar V-gene germline sequence identity to that of the native VL, in order to yield a similar extent of antibody maturation in each shuffled pair, thus making the pair classification task harder, following Ref. (20). In 74% of cases, p-AbNatiV2 assigned a higher pairing likelihood to native pairs than to random pairs (**Fig. 2e**). For comparison, the same assessment was carried out for Humatch and ImmunoMatch (32), which are recently introduced models developed to predict pairing likelihood. For both methods, the score distributions of native and scrambled pairs overlapped substantially (**Supplementary Fig. 14**), and only ∼60% of native pairs were assigned a higher score (**Fig. 2e**).

Although care was taken to generate random pairs that did not overlap with any observed pairs in the dataset, some of these randomly paired sequences may still result in well-folded, correctly assembled and stable Fvs (55). Therefore, we also repeated the evaluations by comparing the score of a native pair with those of 50 random pairs drawn from the same V-gene germline sequence identity bin, to account for the possibility that some of these 50 random pairs may assemble productively, and thus have a pairing likelihood at par or better than that of the native pair. We found that p-AbNatiV2 ranked the native pair within the top 5 (out of 51 pairs) in over 35% of cases (**Fig. 2f**), highlighting its sensitivity to native sequences. In contrast, Humatch and ImmunoMatch ranked the native pair in the top 5 in only 18% and 16% of cases, respectively, compared to a random expectation of 10% (**Fig. 2f)**.

In addition, ImmunoMatch exhibited a characteristic failure mode when evaluating sequences outside of its learned distribution. It assigned high pairing scores to PSSM-generated artificial sequence pairs (**Supplementary Fig. 14b**) and to both paired and unpaired mouse Fv sequences (**Supplementary Fig. 15c**), despite these being distant from the learned distribution of paired human antibodies. In contrast, p-AbNatiV2 and Humatch assigned low pairing likelihood to artificial sequences and to mouse sequences regardless of their native pairing status (**Supplementary Fig. 15a-b**), demonstrating a more interpretable out-of-distribution behaviour. This difference likely stems from the negative sampling strategies used during training. ImmunoMatch was trained exclusively on shuffled and paired sequences (i.e., negatives drawn from the same distribution as the positives). By contrast, p-AbNatiV2 and Humatch incorporate a broader negative class including mismatched sequences from different B-cell classes and studies, with p-AbNatiV2 additionally incorporating cross-species sequences. This broader negative space enables the models to assign high score only to sequences within the learned human paired distribution and low score to out-of-distribution, low-humanness sequences.

Combined with its humanness assessment, which effectively captures contextual information from the other chain, the pairing likelihood prediction head of p-AbNatiV2 provides an accurate explicit prediction for VH/VL pairing, enabling prioritisation of compatible partners for paired-antibody development.

### 3.6 AbNatiV2 for the humanisation of VHHs and Fvs

AbNatiV1 has been previously employed to successfully humanise nanobodies derived from immune libraries or engineered in vitro (2, 14, 27–29). By preserving nanobody nativeness while enhancing humanness, humanised variants are expected to retain favourable biophysical nanobody characteristics while exhibiting improved in vivo behaviour in humans. The humanisation of camelid nanobodies has notably been shown to reduce human T-cell responses, particularly against epitopes in the FR2 and CDR regions (56). Therefore, humanising therapeutic nanobodies has become a rather standard step for therapeutic development, and the majority of clinical candidates have undergone this process (**Supplementary Table 4** and Methods) (1).

AbNatiV2 scoring shows that humanised clinical-stage nanobodies occupy the space that separates human VH and camelid V_H_H sequences in a V_H_H -nativeness versus VH-humanness plot (**Figure 3)**. In comparison, therapeutic antibodies with humanised VH domains span a similar range of humanness scores to humanised V_H_H sequences, showing that successful therapeutic antibodies do not necessarily need to occupy the native space of human sequences (humanness ≥ 0.8, y-axis of **Fig. 3**). For example, the first FDA-approved nanobody drug, Caplacizumab (57), displays a relatively low humanness score of approximately 0.6 (up from the 0.4 of its parental llama-derived V_H_H (58)). We also note that, in contrast with AbNatiV1 (**Supplementary Fig. 16**), the more stringent scoring of AbNatiV2 does not allow for any V_H_H sequence (clinical or not) to score for both humanness and V_H_H -nativeness greater than the 0.8 threshold, revealing inherent trade-offs between the two properties. Taken together, these observations show that AbNatiV2 provides a quantitative framework to study and explore the nativeness and humanness landscape of V_H_H sequences, which is also relevant for de novo design pipelines (13, 59, 60) where nanobody sequences are generated from scratch without direct biological constrains.

**Fig. 3.**
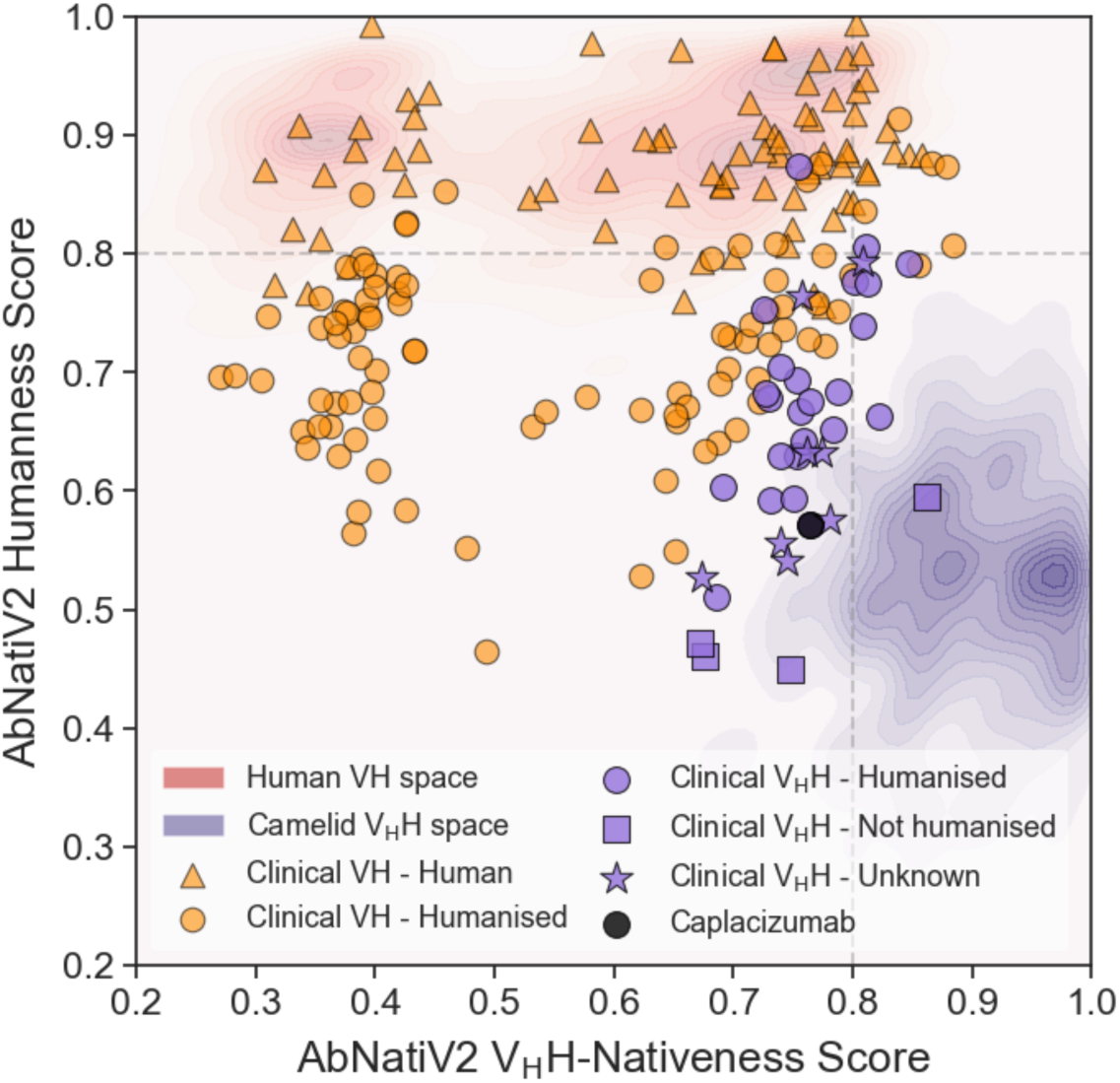
The nativeness landscape of therapeutic antibodies and nanobodies. AbNatiV2 humanness versus V_H_H-nativeness scores visualised as kernel densities for 500 sequences from the human VH test dataset (red areas) and camelid V_H_H test dataset (purple areas). Overlaid are clinical-stage sequences: 170 VH antibodies (orange markers) classified as human (triangles) or humanised (circles) (from Prihoda et al. (53)), and 36 V_H_H nanobodies (purple markers) from the Thera-SAbDab dataset (61) classified as humanised (circles), non-humanised (squares), or unknown (stars) based on manual literature curation (**Supplementary Table 4** and **Methods**). The first FDA-approved nanobody drug, Caplacizumab (57), is in black. The corresponding AbNatiV1 landscape is provided in **Supplementary Figure 16**.

The newly trained AbNatiV2 models are readily applicable for the humanisation of both nanobodies and Fv regions from any source. For nanobodies, the automated pipeline samples humanising mutations that preserve V_H_H-nativeness using the new unpaired AbNatiV2 models (**Supplementary Fig. 17a**). For paired Fvs, the original implementation of AbNatiV1 separately humanised the heavy and light chains. Pairing information was only leveraged at the start, when identifying surface-exposed residues on the modelled structure of the whole Fv. Here we introduce a paired humanisation pipeline (see Methods and **Supplementary Figure 17b**) that jointly optimises the heavy and light chain sequences using p-AbNatiV2. This paired approach scores the heavy and light chains together at each iteration, allowing mutations to be assessed based on their combined effect on both chains. Additionally, pairing likelihood is monitored at each step, and any mutation reducing the initial pairing likelihood beyond a user-defined threshold is rejected. Comprehensive performance characterisation of the paired humanisation approach is left to future work. The paired humanisation pipeline and associated user manuals are available in the AbNatiV2 repository (https://gitlab.developers.cam.ac.uk/ch/sormanni/abnativ).

## 4 Discussion

In this work, we present the unpaired AbNatiV2 and paired p-AbNatiV2 models to perform antibody and nanobody nativeness assessment and engineering. The AbNatiV2 models integrate rotary embeddings, gated attention mechanisms, and a focal loss strategy, resulting in improved performance and accuracy over AbNatiV1, by more effectively capturing high-order relationships between residues and mitigating germline bias common to antibody PLMs (19). The paired p-AbNatiV2 model employs a novel cross-attention-based VQ-VAE architecture equipped with a pairing prediction head. This architecture allows for the simultaneous evaluation of VH/VL humanness and the prediction of their pairing likelihood.

As part of this work, we curated a new dataset comprising 21 million nanobody sequences derived from immune camelid repertoires, with around 9.2 million new sequences coming from deep sequencing experiments we carried out (**Supplementary Table 1**). This datasets marks a substantial increase compared to the largest prior collection, the INDI dataset (37), which contained approximately 11 million sequences. In parallel, we curated a dataset of almost 4 million paired human sequences by combining data from the OAS dataset (17) and the recently published PairedAbNGS dataset (19). These datasets, that we make publicly available, serve as a critical foundation for advancing the development of computational tools for both nanobody and paired antibody engineering.

All AbNatiV1 models (V_H_H, VH, Vκ and Vλ) were each trained on two million sequences. However, the V_H_H corpus had limited diversity, with most sequences drawn from only three studies. This likely accounts for the sharp drop in V_H_H-AbNatiV1 performance when evaluated on the larger, more diverse V_H_H database assembled for AbNatiV2. By contrast, all AbNatiV1 human models did not exhibit a performance decline on the expanded AbNatiV2 human dataset (**Supplementary Figure 8**). Therefore, we suggest that the principal driver of the performance gains in V_H_H -AbNatiV2 model is the expanded and more diverse training set, whereas the (smaller) improvements observed in the unpaired human AbNatiV2 models are primarily attributable to the architectural refinements we introduced.

Capturing VH-VL pairing compatibility is essential for the robust engineering of antibodies and for de novo design applications. By leveraging the newly curated dataset of paired human sequences and incorporating a pairing prediction head, p-AbNatiV2 effectively evaluates the pairing compatibility while addressing its inherent complexity through noise-contrastive learning with both “easy-but- confident” and “hard-but-noisy” negatives. In this framework, true VH–VL pairs are confidently identified as positives, mismatched pairs drawn from different repertoires serve as clear mis-paired examples, and within-batch shuffles act as intermediate “hard-to-spot” negatives yielding a broader pairing likelihood distribution, which reflects that arbitrary pairings of chains from the same repertoire often leads to compatible heavy and light chains (38) (**Fig. 2d**). By coupling partner-conditioned humanness assessment with an explicit VH/VL pairing-likelihood score, p-AbNatiV2 provides an interpretable, ready-to-deploy scoring tool to rank Fv for humanness, prioritise compatible pairs, guide rational mutation design, and complement generative and experimental pipelines.

In summary, AbNatiV2 and p-AbNatiV2 are designed to act as practical scoring layers for antibody design and optimisation pipelines. In generative settings (e.g., diffusion, autoregressive, or inverse- folding models), sequence-level VHH-nativeness or humanness provide an efficient pre- and post-generation filter, while residue-resolved profiles focus mutation proposals, CDR graft choices, and developability edits on positions most compatible with native repertoires. Partner-aware humanness together with an explicit VH/VL pairing-likelihood score enables ranking of candidate light chains for a given heavy chain (and vice versa), supports common light-chain and multispecific formats, and flags variants likely to disrupt pairing or paratope stabilisation. The calibrated scoring (with a consistent operating threshold of 0.8) facilitates multi-objective selection alongside affinity, stability, and manufacturability metrics, and the sequence-only inputs make integration into iterative design-make-test loops straightforward. By releasing models, datasets, and a web server, this work standardises nativeness and pairing assessment as routine criteria, with the expectation that wider adoption will reduce experimental dead-ends and shorten the path to developable leads.

## 5 Methods

### 5.1 Dataset curation

The source and characteristic of each dataset of antibody sequences used in this study is detailed in **Supplementary Table 1**. The unpaired human antibody sequences were downloaded from the Observed Antibody Space (OAS) database (17) in September 2024. The paired human sequences were sourced from the p-IgGen (around 1.6M) (20) and PairedAbNGS (around 2M) (31) datasets. The camelid sequences, spanning alpaca, llama and camel species (respectively, 77%, 16% and 7%; see **Supplementary Figure 4**), were compiled from multiple sources, including the original AbNatiV1 dataset (around 2M sequences) (2), the NGS-sequences from 7 different studies parsed by the Integrated Nanobody Database for Immunoinformatics (INDI) (around 9.4M) (37), a novel dataset derived from the repertoire of 22 non-immunised alpacas (around 660k) (38), and a native non-immune alpaca nanobody library amplified and sequenced for this study (around 9.2M sequences, see after). The additional unpaired mouse and rhesus antibody datasets were sourced from the original AbNatiV1 study, while the paired mouse and rat sequences were downloaded from the paired OAS database.

All datasets will be made available upon publication.

#### 5.1.1 Immune library and MiSeq sequencing

In addition to nanobody sequence datasets sourced from the literature, we produced a library of native alpaca sequences in our laboratory. This non-immune library was constructed as described previously (62, 63). Briefly, peripheral blood mononuclear cells (PBMCs) were isolated from two non-immunised alpacas, and total RNA was extracted using TRIzol reagent (cat#15596026, Invitrogen) according to the manufacturer’s protocol. Protein-coding complementary DNA (cDNA) was synthesised from the total RNA template using oligo dT primers. The variable regions of the VH3 family immunoglobulin gene were subsequently amplified through PCR.

Two separate reactions were performed using the forward primer V_H_H-Lead-F (5’- GTCCTGGCTGCTCTTCTACAAGG-3’) and one of two reverse primers: V_H_H-CH2-R1 (5’- GGTACGTGCTGTTGAACTGTTCC-3’) or V_H_H-CH2-R2 (5’-CGCCATCAAGGTAC- CAGTTGA-3’). Consequently, four individual libraries were constructed, two from each animal. To prevent the carry-over of sequences amplified from conventional immunoglobulin families, rather than heavy-chain-only antibody genes, PCR products around 700 bp were recovered from a 2% TBE agarose gel, while CH1-containing products (900-1000 bp) were excluded.

Next-generation sequencing (NGS) was performed on the Illumina MiSeq platform using a 600-cycle kit to generate 2 × 300 bp paired-end reads. For sequencing library preparation, adapters and barcodes were added to the amplified immune libraries by PCR. To minimise amplification bias toward certain sequences, only 10 PCR cycles were performed. The final libraries were quantified using real-time PCR, normalised to equal molar concentrations, and diluted to 10 nM for the NGS run.

#### 5.1.2 NGS data analysis

NGS data were obtained from the MiSeq sequencing platform as two fastq files, one for forward and one for reverse reads. These files were processed with an in-house python script that merges forward and reverse reads assuming an overlapping region (which is always present for short proteins like V_H_Hs) to obtain the full sequence of the V_H_Hs. The script further counts the number of times each unique sequence is observed in the raw files (i.e., the number of reads per unique sequence) and uses phred scores to estimate the probability of observing at least one nucleotide sequencing error in the nanobody domain. Sequences were in silico translated to amino acids. All V_H_H sequences observed at least 2 times and with a probability of containing at least one nucleotide sequencing error lower or equal to 0.001 were kept.

#### 5.1.3 Sequence alignment and cleaning

All antibody sequences were aligned with the ANARCI software (64) using the AHo numbering scheme (45). Sequences not containing conserved cysteines and with partial C- or N-terms were discarded, as described previously in AbNatiV1 (2). AbNatiV1 was always run with its own AHo- alignment protocol, but for AbNatiV2 minor adjustments were made, when necessary, to ensure that each CDR contains a single and continuous gap region, as expected by the AHo numbering (45). We found that ANARCI running with the AHo numbering scheme often inserted an isolated gap at position 28 of the CDR1 stem, detached from the main gap region expected at the centre of the loop (45). We found that this isolated gap can spread to positions 27 and 26 in Kappa chains (2) and alter the number of residues included in the CDR1, which is defined to begin at position 27. To address this inconsistency, we merged all gaps found between the conserved cysteine at position 23 and the end of the CDR1 into one gap region centred in the middle of the CDR1. We applied the same protocol to CDR2 and CDR3 loops.

We also observed in the original camelid V_H_H dataset of AbNatiV1 that a histidine was present at the N-terminus in 21% of the sequences (**Supplementary Fig. 18**). Yet, this histidine prevalence was not observed in our library of native V_H_Hs (**Supplementary Fig. 3**) and is similarly not observed in the germline genes annotated onto IMGT (65). This high prevalence of N-terminal histidine in the AbNatiV1 V_H_H dataset is therefore likely resulting from commonly used degenerate amplification/sequencing primers (typically starting with the DNA sequence ‘SAKGTG…’), which is designed to accommodate the naturally occurring EV/QV/DV variation in the amino acid sequence at the beginning of the FR1 region, but that also encodes for HV (66). Conversely, our sequenced library was amplified from the conserved leader signal sequence. For instance, Xiang *et al.*, whose sequenced camelid repertoire was a major source of sequences for the AbNatiV1 training set, used a degenerate primer containing the 5’-SAK-3’ motif, likely leading to an artificial abundance of N-terminal histidine. Consequently, we replaced every N-terminal histidine with either a glutamine, a glutamic acid, or an aspartic acid at random. This N-terminal histidine cleaning was only applied to the AbNatiV1 and INDI datasets, as they both contain sequencing studies employing degenerative primers on the N-terminus of the variable domain. Before processing, they exhibited respectively 21% and 3% of N-terminal histidine. Finally, all camelid datasets were further filtered using AbNatiV1, removing sequences with an AbNatiV1 V_H_H-ness score below 0.4, to eliminate sequences that are extremely far from the learnt distribution (even further away than human VH or artificial PSSM-generated sequences are (2)), and hence likely contain amplification, assembly, or sequencing errors.

For the human antibody datasets, only sequences appearing at least twice in the original OAS database, which contains approximately 2 billion human sequences, were selected to minimise the use of sequences affected by sequencing errors.

#### 5.1.4 Parsed unpaired datasets

After curation, the final datasets contain 19,654,973 human VH, 21,228,567 human VLight (including around 50% of kappa chains and 50% of lambda chains), and 20,208,125 camelid V_H_H unique sequences. Each AHo-aligned dataset was clustered with the Linclust algorithm (67), using a 95% sequence identity threshold on target coverage (i.e., coverage mode 1). 100,000 representative sequences from different clusters were selected for validation, and 50,000 for testing. The remaining sequences were used for training. The VKappa and VLambda sequences were combined to train a single light model into one dataset, named VLight dataset.

The test set built in this way is representative of the broad sequence distribution but likely contains sequences relatively similar to those in the training set. Therefore, to evaluate the ability of AbNatiV2 to generalise to more distant sequences, an additional diverse test set was generated for each model. Diverse sequences are at least 2.5% distant from any sequence in the corresponding training set of the human VH and VLight datasets, and 5% for the camelid nanobody training set. A lower divergence threshold for human VH and VL was required because too few sequences were available at 5% divergence from their closest training-set sequence to support a robust evaluation. By contrast, a 5% threshold remained feasible for nanobodies, likely because VHH repertoires are more diverse than human VH or VL considered individually, and because the VHH dataset pools sequences from multiple camelid species further increasing diversity. The 5% setting provides a more stringent test of generalisability. Distance was defined as the edit distance normalised by the gapless length of the test sequence under scrutiny. The first 10,000 sequences identified were selected to form the diverse test set (**Supplementary Fig. 19**).

The residue frequency matrices (or position weight matrices; PWMs) and corresponding position-specific scoring matrices (PSSMs) of human and camelid datasets are presented in **Supplementary Figure 2**. For each model, a dataset of 10,000 artificial sequences, named PSSM-generated, was generated filling each sequence position by randomly picking residues using the residue frequency observed at that position. Sequences were re-aligned to prevent multiple gaps in CDR regions, which may be easily identifiable by the trained models and bias the classification task. For the VLight model 5,000 VKappa and VLambda sequences were separately generated from their corresponding PWM, which roughly corresponds to the ratio of VKappa and VLambda sequences in the VLight dataset.

#### 5.1.5 Parsed paired datasets

After processing, the final paired dataset contained 3,745,614 unique paired human sequences. The paired sequences were concatenated to be clustered with Linclust, as previously described. 100,000 representative sequences from different clusters were selected for validation, and 50,000 as test set, ensuring that none were included in the training datasets of the AbNAtiV2 unpaired VH and VLight models. The diverse test set contains sequences at least 5% distant from any sequence in the training set.

3,745,614 more mismatched sequences were obtained from Humatch (21), comprising mixed heavy and light chains from different B-cell classes (e.g., memory and naïve) and OAS studies. Alongside those sequences, we included as well 119,311 shuffled mouse sequences derived from mouse sequences in the PairedAbNGS dataset (31). Furthermore, each VH mouse sequence was artificially paired with a random VL human sequence from the human dataset, and reciprocally with VL mouse sequences. We employ these mismatched sequences as negative examples in the noise-contrastive learning.

### 5.2 The unpaired AbNatiV2 model

#### 5.2.1 Unpaired architecture

AbNatiV2 is a vector-quantised variational auto-encoder (VQ-VAE) built with transformer blocks in a latent space compressed by convolutional patches (see **Figure 1a**). Compared to AbNatiV1 (see more details in Ref. (2)), the new model presents significant architectural changes (**Supplementary Figure 20**). The sinusoidal positional encoding in the embedding layer is replaced with rotary positional embeddings within the transformer layers, facilitating the encoding of relative positional information (40). The output of the scaled dot-product attention mechanism is now dynamically modulated by an integrated gating mechanism in each layer (41). The multi-layer perceptron (MLP) of the residual connection is replaced by a SwiGLU activation mechanism, which improves efficiency and representation capacity (42).

All the hyperparameters were manually tuned for the V_H_H model, effectively resulting in an up-scaling of the AbNatiV1 model, as expected from the employment of a ∼10x larger training set (**Supplementary Table 2**). Specifically, the number of layers was increased from 3 to 8, the number of attention heads from 8 to 16, and the embedding dimension from 768 to 1,024. Since we observed that the hyperparameters of the AbNatiV1 did not significantly vary between camelid and human training sets of comparable size, the same hyperparameter values were used also for all AbNatiV2 human models. Training was stopped upon convergence (see **Supplementary Figure 5**). The V_H_H model was trained for 30 epochs, while the VH and VLight models were trained for 35 epochs.

#### 5.2.2 Loss function of the unpaired models

AbNatiV2 unpaired models are trained to minimise a negative evidence lower bound (NELBO) composed of a reconstruction term and a vector quantisation commitment term. A focal reconstruction loss (FRL) based on mean-squared error (MSE) is used for the reconstruction term:

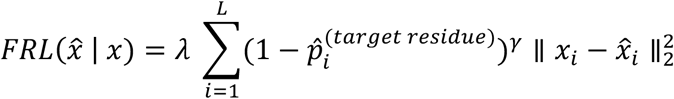

The FRL of the output *x̂*, given the training input sequence 𝑥 of length 𝐿, is a weighted summation of the positional MSEs. Each positional MSE is modulated by 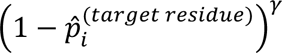, where 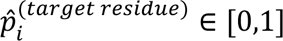 represents the output likelihood probability assigned to the target residue (i.e., the residue present in the training sequence at position *i*) in *x̂* at position 𝑖. The parameters λ and γ serve respectively as linear and quadratic scaling factors, with λ = 1 and γ = 1 providing the best performance. In the FRL, positions that the model reconstructs with high accuracy (i.e., when 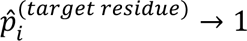) are dynamically down weighted at each step. It reduces the loss contribution of positions where the model is confident, such as for conserved residues in the framework regions, driving the model to allocate more training resources (i.e., weights) to more variable regions.

A static version of the FRL (sFRL) is also implemented by directly weighting the positional MSE at each position with the corresponding conservation index computed from the PSSM of the training dataset (**Supplementary Fig. 3**) using the variance-based approach described in Ref. (68). The sFRL is defined as:

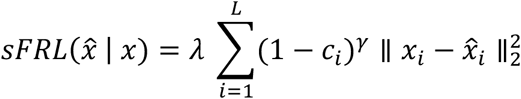

Here, the dynamic 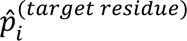 of the FRL is replaced by 𝑐_𝑖_ ∈ [0,1], the pre-computed conservation index at position 𝑖. When a position is highly conserved (i.e., the same residue very frequently appears at that position), 𝑐_𝑖_ → 1, reducing its loss contribution. This static approach explicitly down-weights the contribution of highly conserved positions.

The neural network was implemented using PyTorch.2.2 (69) and enhanced by the PyTorchLightning.2.4 module (70). The models were trained with a batch size of 128 by the Adam optimizer (71) with a learning rate of 2e-05. During training, the same residue masking strategy of AbNatiV1 was applied to the one-hot encoded inputs.

#### 5.2.3 Antibody nativeness definition of unpaired sequences

The definition of antibody nativeness in AbNatiV2 remains consistent with that in AbNatiV1:

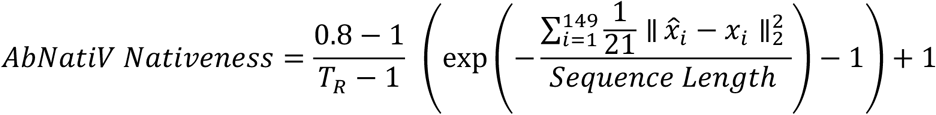

where 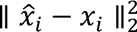 is the MSE at sequence position 𝑖 between the aligned input sequence 𝑥 and the reconstructed output sequence *x̂* reconstructed by AbNatiV. The exponential transformation of the normalised MSE, without focal weighing, is linearly rescaled for interpretability using updated optimal thresholds 𝑇_𝑅_, specific to each trained model (𝑇_𝑅_(𝑉𝐻) = 0.994, 𝑇_𝑅_(𝑉𝐿𝑖𝑔ℎ𝑡) = 0.996, 𝑇_𝑅_(𝑉𝐻𝐻) = 0.993; see **Supplementary Fig. 21**). 𝑇_𝑅_represents the optimal threshold that best separates native sequences (e.g., human or camelid) from non-native sequences (e.g., mouse).

### 5.3 The paired AbNatiV2 model

#### 5.3.1 Paired architecture

The pre-trained AbNatiV2 VH and VL unpaired models were combined using cross-attention and noise-contrastive learning to develop a paired VH/VL model, named p-AbNatiV2, that jointly processes paired antibody sequences and predicts both humanness and pairing likelihood of a given pair of VH and VL sequences (**Fig. 2c**). p-AbNatiV2 is composed of two units: a cross-attention VQ-VAE which projects and reconstructs heavy and light chain jointly, and a pairing prediction head which classifies paired sequences versus mis-paired ones.

In the cross-attention VQ-VAE, the two unpaired architectures are interconnected through cross-attention layers (52, 72, 73) integrated within the transformer blocks of both the encoders and decoders. Specifically, between two transformer layers of the heavy encoder, the output of the preceding transformer layer serves as the query for a cross-attention layer, while the corresponding output from the light model as the key and value in the scaled dot product operation. The resulting output from the cross-attention layer is then given back to the heavy model as input to the subsequent transformer layer. This operation is symmetrically mirrored for the light transformers and then repeated in the decoder, post discrete vectorisation and their respective heavy and light codebooks. The pre-trained weights of the unpaired models were kept constant and only the weights of the cross-attention layers were updated during training. The transformer layers for cross-attention share the same architecture as those in the unpaired models, except that the sigmoid cross-gating mechanism is initialised with a low bias term of −5 to facilitate the integration of the cross information in the unpaired models, as too much signal coming from the still untrained cross-attention weights would likely disrupt the reconstruction task early in training (73).

In the pairing prediction head, the heavy and light outputs from the final cross-attention layer, prior to deconvolution in the decoder, are combined using a dot product following a linear transformation. The resulting output is passed through a sigmoid-linear projection head, which predicts the pairing compatibility between given VH and VL inputs (i.e., the likelihood that the input heavy and light chains form a native pair). The LR model is trained using noise-contrastive learning (54), where native sequence combinations from the training set are labelled as positive ‘paired,’ while negative ‘mis- paired’ are generated by either shuffling the light chains within the current training batch, or by mismatching heavy and light chains from different human B-cell classes, species and studies. Including some non-human training sequences is important; otherwise, this prediction head would see only high- humanness sequences as input and the pairing likelihood would be unlikely to generalise to low- humanness inputs. Such inclusion reduces the risk of assigning spuriously high scores to low- humanness sequences, but it does not make the model suitable for predicting pairing likelihoods of antibodies from other species.

Because we employed two distinct negative examples (i.e., shuffled and mismatched), we defined a weighted binary cross entropy, 𝐵𝐶𝐸_5𝑒𝑖gℎ𝑡𝑒𝑑_, to balance their contributions as follows:

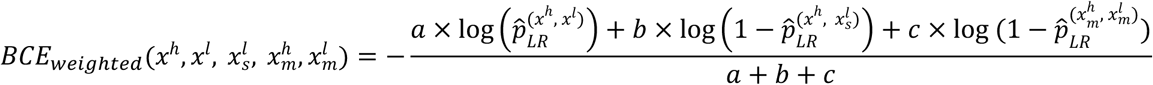

Here, 𝑥^ℎ^ and 𝑥^𝑙^ represent the input heavy and light chain sequences. 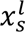 corresponds to the shuffled light chain in the negative shuffled example, and 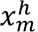, 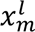 the mismatched negative example. *p̂_LR_* is the predicted pairing label computed by the LR head model for a given pair. 𝑎, 𝑏 and 𝑐 correspond to the corresponding loss weight associated with each example. We manually finetuned these weights, ultimately selecting 𝑎 = 1 for native pairs, 𝑏 = 0.5 for shuffled pairs, and 𝑐 = 0.5 for mismatched pairs (**Supplementary Figure 12**).

Residue masking is applied to each heavy and light sequence throughout the training process. All hyperparameters were manually tuned (**Supplementary Table 2**). The paired model was trained for 20 epochs.

#### 5.3.2 Paired loss

The p-AbNAtiV2 model is trained using the paired loss (𝑝𝐿), defined as:

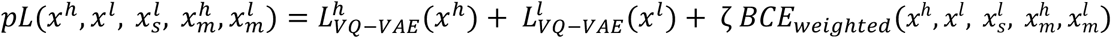

Here, 𝑥^ℎ^ and 𝑥^𝑙^ represent the input heavy and light chain sequences. 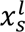 corresponds to the shuffled light chain in the negative shuffled example, and 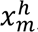, 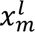 the mismatched negative example. 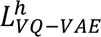 and 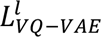 are the respective losses of the heavy and light models, which are backpropagated through the cross-attention layers. 𝐵𝐶𝐸_w𝑒𝑖gℎ𝑡𝑒𝑑_ the binary multi-negative cross-entropy loss, which evaluates the prediction error of the pairing head. ζ = 0.005 is a scaling factor that weights the contribution of the contrastive loss component.

#### 5.3.3 Antibody paired humanness and pairing likelihood definitions

For a given pair of heavy and light chain sequences, p-AbNatiV2 return three humanness scores: one computed on the concatenated VH and VL MSE, one based solely on the VH, and another only for the VL. As for the unpaired models, each score is linearly rescaled using its specific 𝑇_𝑅_ (𝑇_𝑅_(𝐹𝑣) = 0.993, 𝑇_𝑅_(𝑉𝐻𝑝𝑎𝑖𝑟𝑒𝑑) = 0.992, 𝑇_𝑅_(𝑉𝐿𝑝𝑎𝑖𝑟𝑒𝑑) = 0.995; see **Supplementary Fig. 21**), so that 0.8 is consistently the best threshold to separate human from non-human sequences. In addition, p- AbNAtiV2 returns 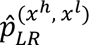 which quantifies the pairing likelihood between the input VH and VL sequences.

### 5.4 Performance metrics

All the reported performance metrics were computed by analysing 10,000 scored sequences for each database. For datasets smaller than 10,000, the whole dataset was used. Precision-recall (PR) classification tasks and amino acid reconstruction accuracy were calculated following the same procedure of AbNatiV1. The benchmarking mouse and rhesus unpaired datasets are new datasets extracted from new expended repertoires from the OAS for mouse, and from the OAS and new NGS data from Ref. (74) for rhesus.

#### 5.4.1 Predictions on antibody therapeutics

In this study, the IMGT dataset (53, 75) includes 532 paired therapeutics. By comparison, AbNatiV1 included 549 therapeutics as parsed by BioPhi (53). Here, only the therapeutics with a single heavy chain paired with a single light chain were chosen, ensuring compatibility with the paired model. Of these, 342 were classified as non-human (humanized, chimeric, or mouse), and 190 as fully human.

The immunogenicity dataset contains 216 paired therapeutics with an anti-drug-antibody (ADA) response (i.e., the percentage of patients who developed ADAs during clinical trials). From the original dataset, as parsed by BioPhi (53), Pexelizumab was discarded due to a missing C-terminus that prevented alignment.

When using unpaired models, the AbNatiV2 humanness score of a paired set of sequences was computed as the average of VH and VL scores. In addition to the other humanness assessment previously used for benchmarks in AbNatiV1 (53, 76–80), we compared AbNatiV2 with the perplexity scores of the paired therapeutic sequences computed by p-IgGen (20) and Humatch (76).

#### 5.4.2 Grafting assessment on nanobodies

The experimental grafting dataset from AbNatiV1 was used. It contains the binding affinity (K_D_) of 6 nanobodies and their variants grafted onto the camelid universal framework (UF), as measured in Ref. (47). Additionally, a dataset of in silico grafted variants was generated by replacing the CDRs of the UF scaffold with those from 10,000 different nanobodies from the new camelid test set, used to assess both AbNatiV1 and AbNatiV2 models.

#### 5.4.3 Pairing likelihood prediction

The ability of p-AbNatiV2 to predict pairing likelihood is quantified by its capacity to distinguish a test-set VH sequence paired with its native VL from the same VH paired with random VL or mismatched VL. Performance was evaluated using the likelihood from the pairing prediction head.

For 1 vs 1 comparisons, 10,000 mis-paired VH/VL were generated by randomly scrambling the VL of 10,000 VH/VL paired test sequences. For each VH, the new shuffled VL had to belong to the same V-gene germline identity bin as its native VL, following the approach in Ref. (20). The bins were defined as [100–95[, [95–90[, [90–80[, and [80–0] percent sequence identity to germline.

All new combinations were verified to not exist in the original VH/VL paired human dataset, ensuring that natively paired sequences were never erroneously classified as mis-paired negatives.

The same study was employed on the diverse paired test set by shuffling the light chains together (see **Supplementary Fig. 13**). No overfitting was observed since p-AbNatiV2 performs similarly well on the diverse pairs.

#### 5.4.4 Comparing pairing performance of p-AbNatiV2 with Humatch and ImmunoMatch

When comparing pairing performances with Humatch (21), sequences were scored via the line ‘Humatch-classify’ command-line function as provided in their GitHub repository (https://github.com/oxpig/Humatch). The Humatch ‘CNN_P’ scores were directly used to assess pairing compatibility, as defined in their original publication (**Supplementary Fig. 14a**).

When comparing pairing performances with ImmunoMatch (32), the light chain isotype (i.e., kappa or lambda) of each tested sequence was first determined by ANARCI (64) and the corresponding pairing score was computed with the respective ImmunoMatch-κ and ImmunoMatch-λ models (**Supplementary Fig. 14b**) using the functions provided in the example notebook available in their GitHub repository (https://github.com/Fraternalilab/ImmunoMatch/).

We randomly scrambled the VH and VL sequences from the 6,179 paired mouse Fvs of the paired OAS dataset used in **Supplementary Table 3** to compare pairing assessments of all three models on different species (**Supplementary Fig. 15**).

#### 5.4.5 Predictions on clinical-stage nanobodies

We mapped the nativeness landscape (**Fig. 3**) of clinical-stage nanobodies using 36 therapeutic nanobody sequences curated and extracted from Thera-SAbDab (61) by Gordon et al. in the Therapeutic Nanobody Profiler (81). The humanisation status of each sequence (i.e., whether the sequence was humanised or not) was determined through manual review of the original literature, including clinical trial reports and primary research publications describing each therapeutic candidate. The curated dataset along with source references is provided in **Supplementary Table 4**.

### 5.5 Implementation of the Fv humanisation with p-AbNatiV2

An implementation of the humanisation pipelines with AbNatiV2 is made available in its repository (https://gitlab.developers.cam.ac.uk/ch/sormanni/abnativ).

For V_H_H humanisation with the unpaired AbNatiV2 models (**Supplementary Fig. 17a**), the WT nanobody sequence is first modelled using NbForge (82) to identify surface-exposed residues for mutation. Non-CDR surface residues with low AbNatiV2 humanness are selected for humanisation. Humanising mutations are sampled while maintaining V_H_H-nativeness above a user-defined threshold relative to the WT sequence. Ultimately, the humanised variant is modelled, structurally superimposed to the WT model using only the framework, and CDR displacement is monitored by RMSD to assess the quality of the humanisation.

For Fv humanisation with the p-AbNatiV2 model (**Supplementary Fig. 17b**), the pipeline jointly optimises the heavy and light chain sequences. To do so, it differs from the unpaired approach in three key aspects. First, the paired p-AbNatiV2 model scores the heavy and light chains together at each iteration, allowing mutations to be assessed based on their combined impact on both chains. Second, the initial sorting of positions by their sensitivity to other mutations – a step to improve mutational search convergence in the Enhanced pipeline as described in Ref. (2) – is performed simultaneously on both chains. Third, to preserve pairing integrity, pairing compatibility is monitored at each iteration and mutations are rejected if they result in a decrease beyond a user-defined threshold relative to the initial pairing likelihood. The structural modelling steps are the same as the nanobody pipeline, but using AbodyBuilder3 (83) to model the Fv region. To improve the robustness of residue selection, the solvent-accessible surface area of the protein is averaged over 10 predicted structures across 10 different seeds.

## 6 Code and data availability

The AbNatiV2 source code including the new trained models and paired humanisation pipeline are available at https://gitlab.developers.cam.ac.uk/ch/sormanni/abnativ. A user-friendly webserver to run AbNatiV2 is provided at www-cohsoftware.ch.cam.ac.uk/index.php/abnativ. To access the webserver, users need to register a free account and log in. The updated codebase remains fully compatible with the original AbNatiV1 models.

All the parsed training, validation and testing datasets employed in this study are available online at https://zenodo.org/records/17466150.

## 7 Author contributions

AR and PS conceived the project. PS supervised the project. AR developed the deep learning architecture with the guidance of MG and PS. AR parsed and collected the training data with the help of NF and HZ. XX and SO generated the nanobody MiSeq data. AR and PS critically analysed the results and wrote the first version of the paper. All authors analysed data and edited the paper.

## 8 Acknowledgments

P.S. is a Royal Society University Research Fellow (grant no. URF\R\251013). We acknowledge funding from UK Research and Innovation (UKRI) Engineering and Physical Sciences Research Council (EPSRC grant no. EP/X024733/1, an ERC starting grant to P. S. underwritten by UKRI). M.G. is a Yusuf Hamied Graduate Scholar.

