## Supplementary material for "Deep learning assessment of nativeness and pairing likelihood for antibody and nanobody design with AbNatiV2": SI

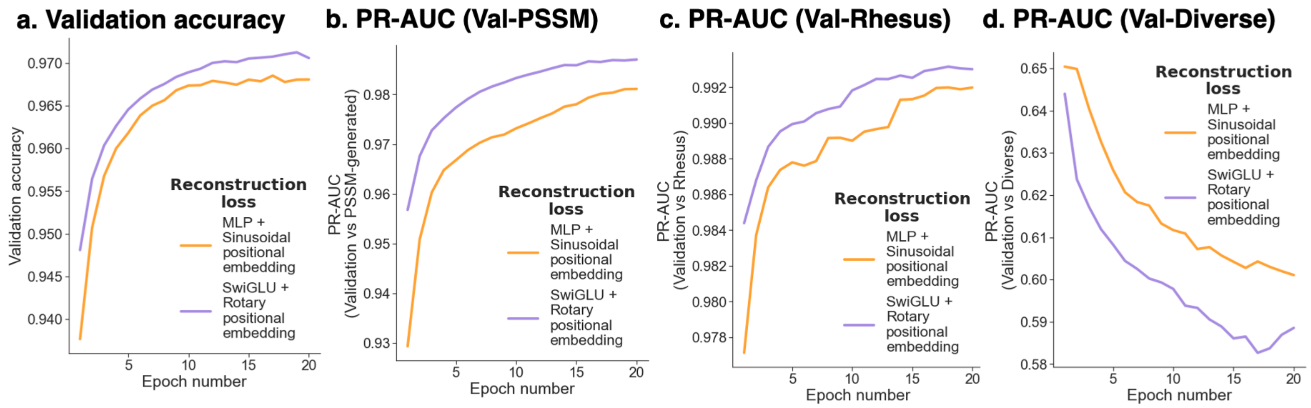


**Supplementary Fig. 1. Impact of the SwiGLU activation function and rotary positional embedding on performance**. The models were trained on the same $V_HH$ dataset with the same hyper-parameters while replacing the multi-layer perceptron (MLP) in the transformer layers' residual connections and the sinusoidal positional embedding (in orange, as employed in the original implementation of AbNatiV) with, respectively, a SwiGLU activation mechanism and a rotary positional embedding (in purple, as in AbNatiV2). (**a**) Amino acid reconstruction accuracy on the Camelid $V_HH$ validation set as a function of training epoch. (**b**) PR-AUC between the nanobody nativeness scores of Camelid $V_HH$ validation set and the $V_HH$ PSSM-generated one. (**c**) PR-AUC between the nanobody nativeness scores of the Camelid $V_HH$ validation set and the VH Rhesus one. (**d**) PR-AUC between the nanobody nativeness scores of the Camelid $V_HH$ validation set and the $V_HH$ diverse test one. As both are datasets of camelid sequences, a PR-AUC closer to 0.5 indicates better performance. The hyperparameters used were identical to those in the final AbNatiV2 model. PR curves were generated using 10,000 sequences per dataset.

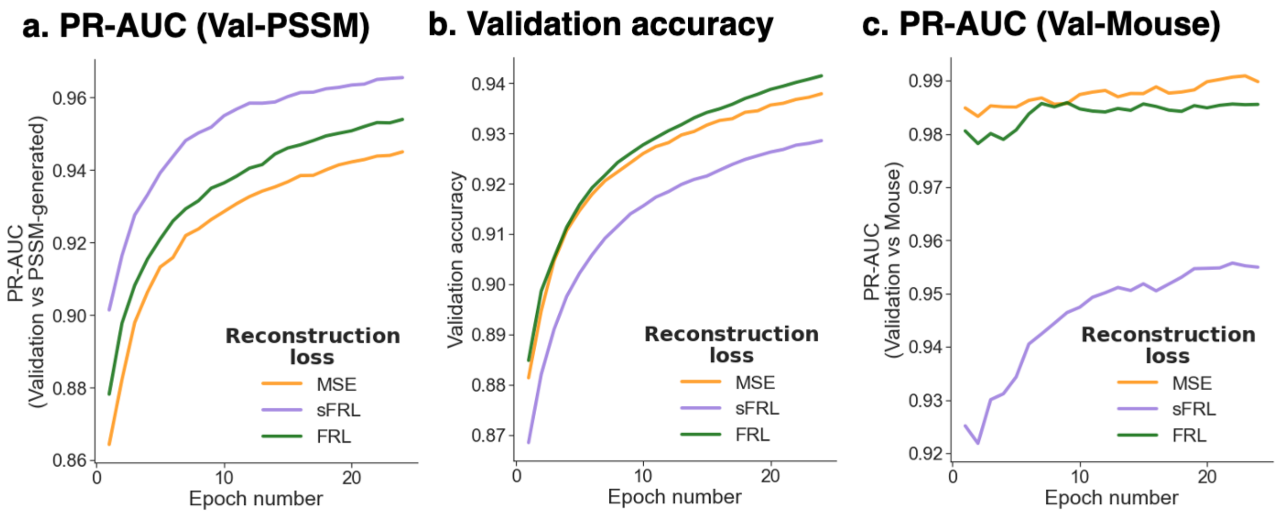


**Supplementary Fig. 2. Impact of the reconstruction loss type on performance**. The models were trained on the Camelid $V_HH$ dataset with the same hyperparameters and architecture of AbNatiV1 while changing the reconstruction loss type: focal reconstruction loss (FRL, in green), static FRL (sFRL, in purple), or standard mean-squared error (MSE, in orange), as employed in the original implementation of AbNatiV. The sFRL approach replaces dynamic focal down-weighting in FRL with a residue-position conservation index penalty. (**a**) PR-AUC between the nanobody nativeness scores of the Camelid $V_HH$ validation set and the $V_HH$ PSSM-generated one across training epoch. (**b**) Amino acid reconstruction accuracy on the Camelid $V_HH$ validation set. (**c**) PR-AUC between the nanobody nativeness scores of the Camelid $V_HH$ validation set and the VH Mouse one. PR curves were generated using 10,000 sequences.

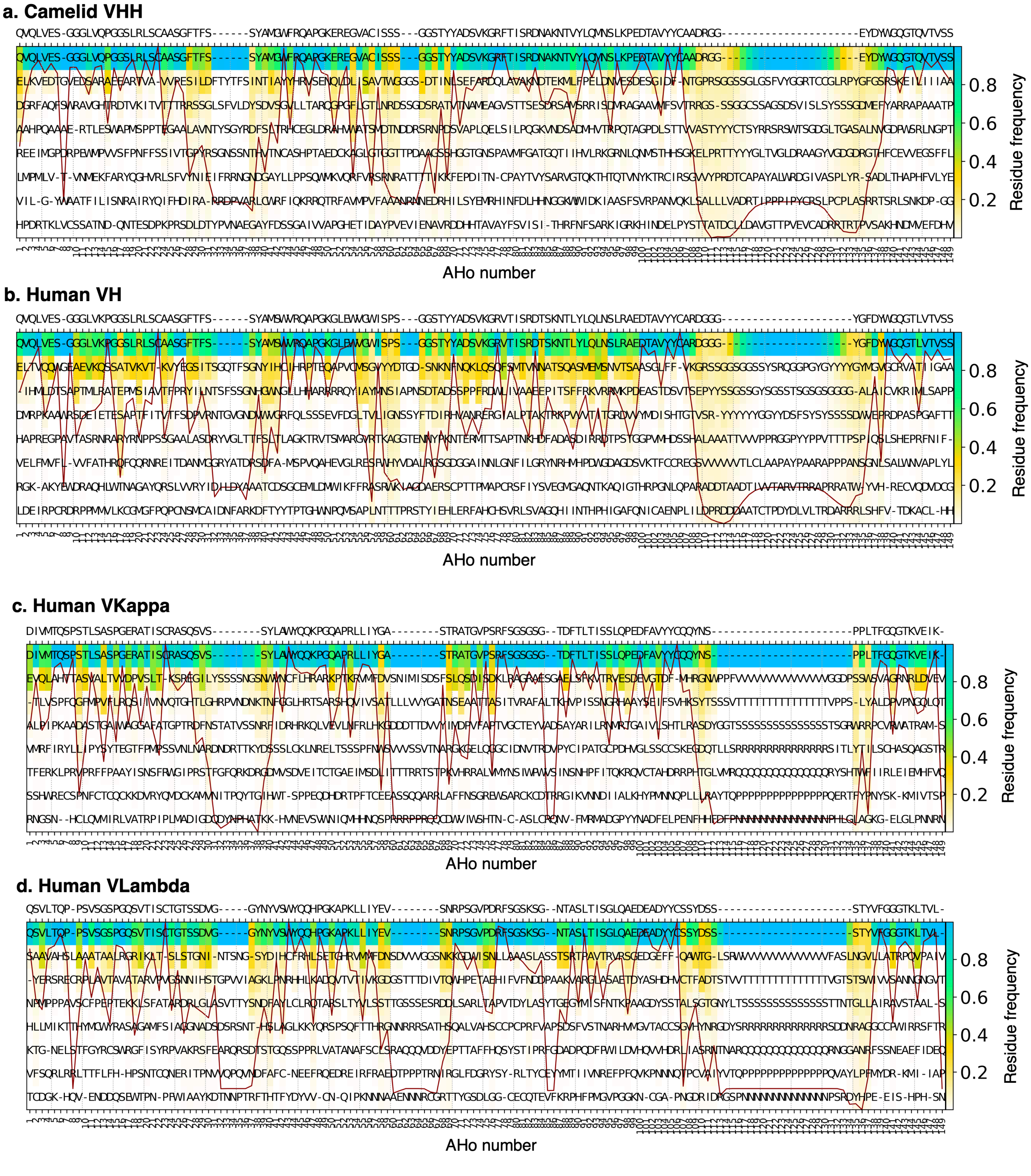


**Supplementary Fig. 3. Position-specific weight matrix (PWM) of the training datasets of AbNatiV2**. PWMs are calculated respectively on the test set of the Camelid $V_HH$ (**a**), Human VH (**b**), Human VKappa (**c**), and Human VLambda (**d**). The x-axis reports the AHo numbering scheme used for the alignment, the colour-bar is the amino acid frequency at each position. Each column is sorted from most to least frequent residue at that position, and only the top eight residues are shown. Residues are coloured based on their frequency at each position (see colour-bar). The consensus sequence, which consists in the most frequent residue at each position, is shown on the top. The continuous red line is the conservation index of each position (high means position highly conserved, low poorly conserved).

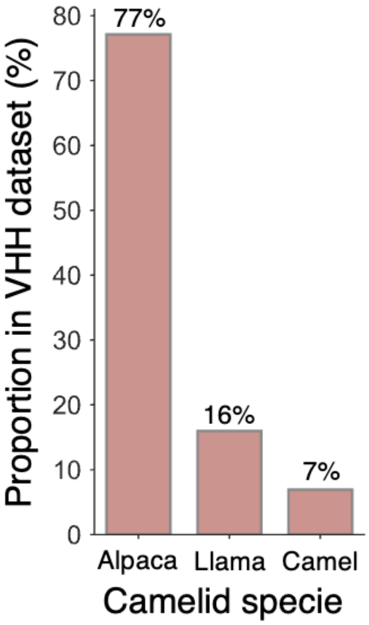


**Supplementary Fig. 4. Distribution of camelid species across the Camelid $V_H$H dataset of AbNatiV2**. The camelid species were retrieved from the original studies (see Methods for the complete list of sources). The proportion for each specie was computed on the compiled pre-processed dataset (before alignment, cleaning, and filtering for unique sequences).

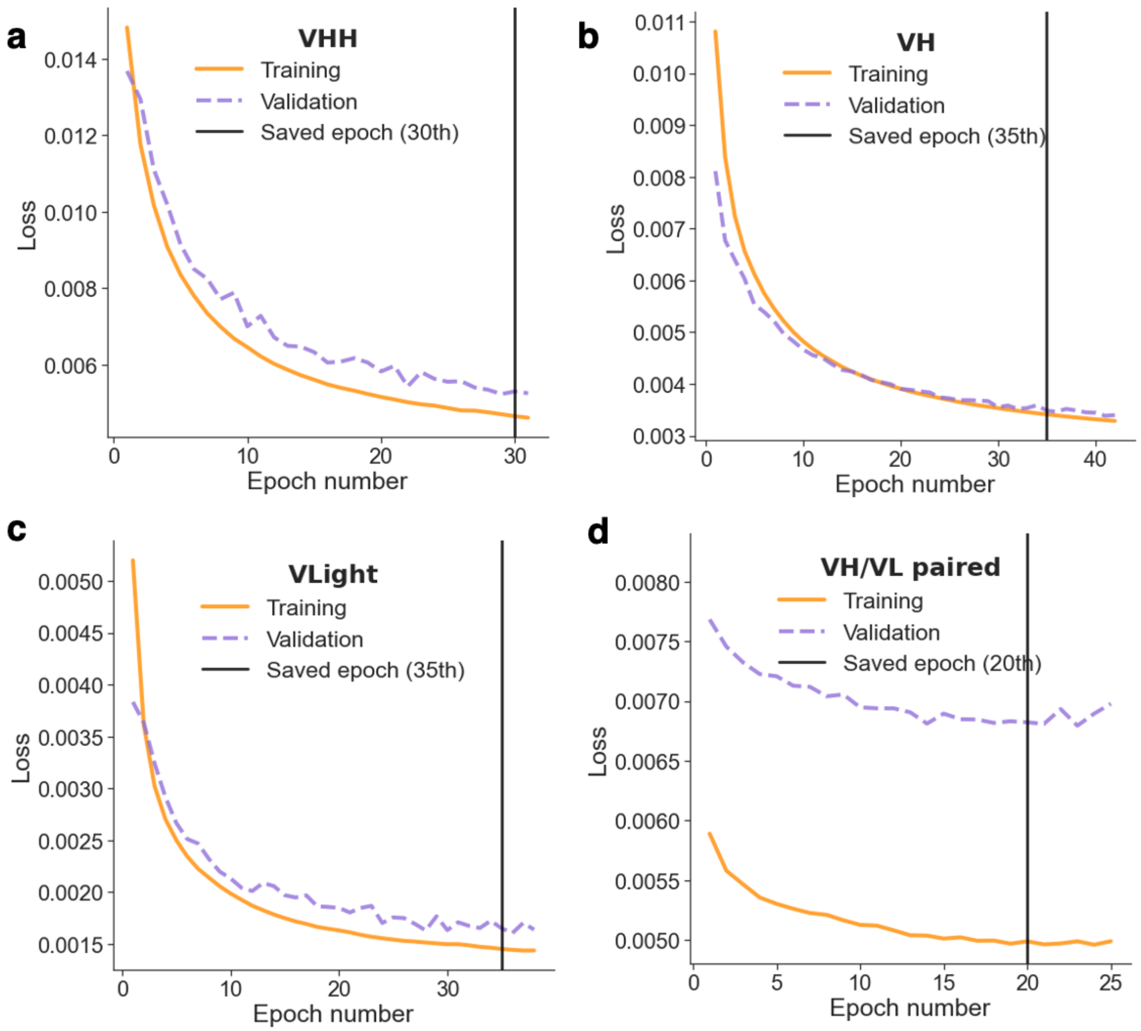


**Supplementary Fig. 5. AbNatiV2 training performance.** Loss function averaged on the whole training set (orange solid line) and validation set (dotted purple line) at each training epoch of the AbNatiV2 model train on camelid $V_HH$ (**a**), human VH sequences (**b**), human VL sequences (**c**), and human VH/VL paired sequences (**d**). The VH and VL models were trained for 35 epochs, and the $V_HH$ model for 30 epochs. The VH/VL paired model was trained for 20 epochs. The black vertical line represents the selected epoch saved for each model and used in AbNatiV2. For the paired model, the loss adds up the VQ-VAE loss and the pairing prediction accuracy. Masking was not applied at the validation stage.

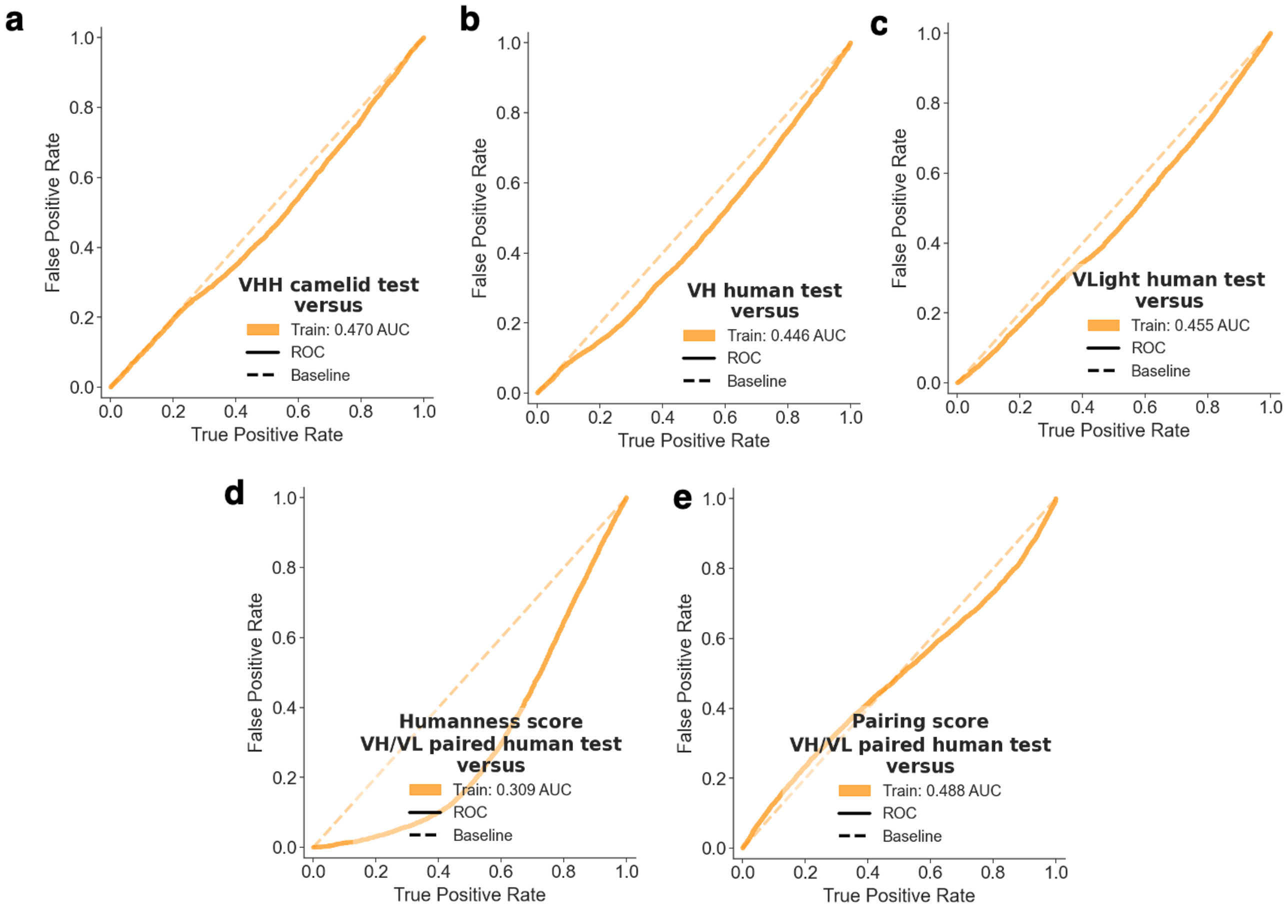


**Supplementary Fig. 6. Overfitting assessment of the AbNatiV models**. ROC curves of the ability of AbNatiV to differentiate the humanness score distribution of the Training dataset (random subset of 10,000 sequences) and the Test dataset (also 10,000 sequences). The analysis is carried out on (**a**) camelid $V_H$H sequences, (**b**) on human VH sequences, (**c**) on human VLight sequences, and (**d**) on human VH/VL paired sequences, always using the corresponding AbNatiV2 model (and p-AbNatiV2 for **d**). In (**e**) the ROC curve was computed using the score from the pairing prediction head of the p-AbNatiV2 model. An AUC of 0.50 corresponds to perfectly overlapping distribution.

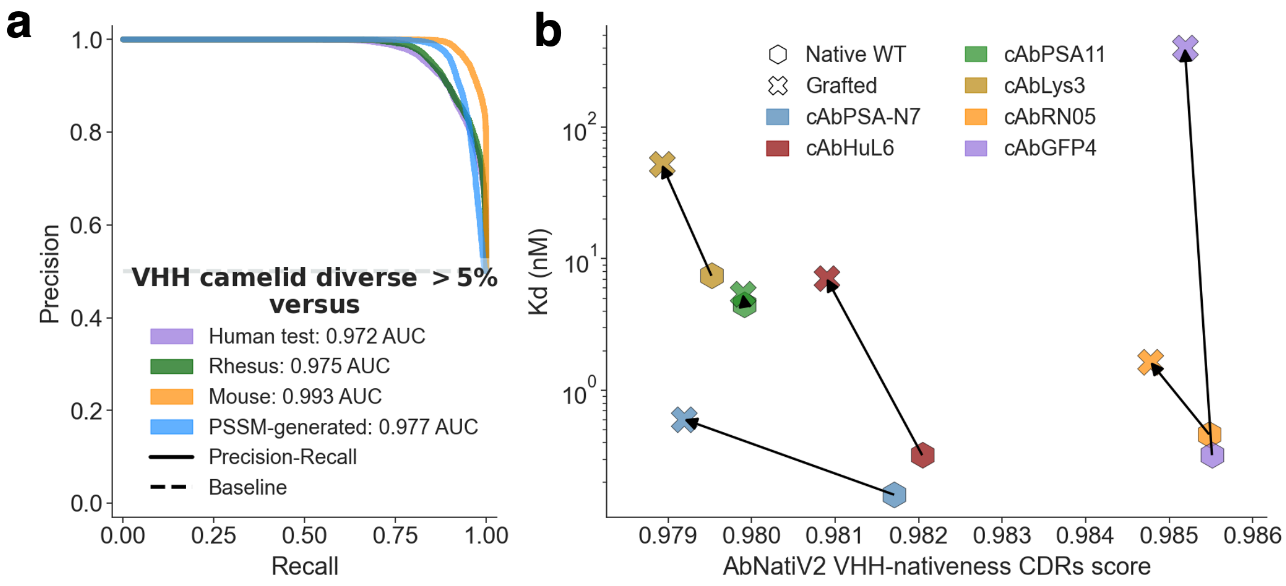


**Supplementary Fig. 7. Additional performances of the AbNatiV2 $V_H$H model**. (**a**) Plots of the PR curves used to quantify the ability of AbNatiV2 to distinguish the $V_H$H camelid diverse test set from the other datasets (see legend, which also reports the AUC values). The baseline (dashed line) corresponds to the performance of a random classifier. (**b**) Plot of the binding $K_D$ (as reported in Saerens *et al.,* J.M. Biol. 2005) as a function of the AbNatiV2-$V_H$H nativeness score averaged across all CDR position of six nanobodies (legend) before and after grafting of all three CDRs onto a camelid UF. An arrow is directed from the native sequence in the WT framework to the grafted one.

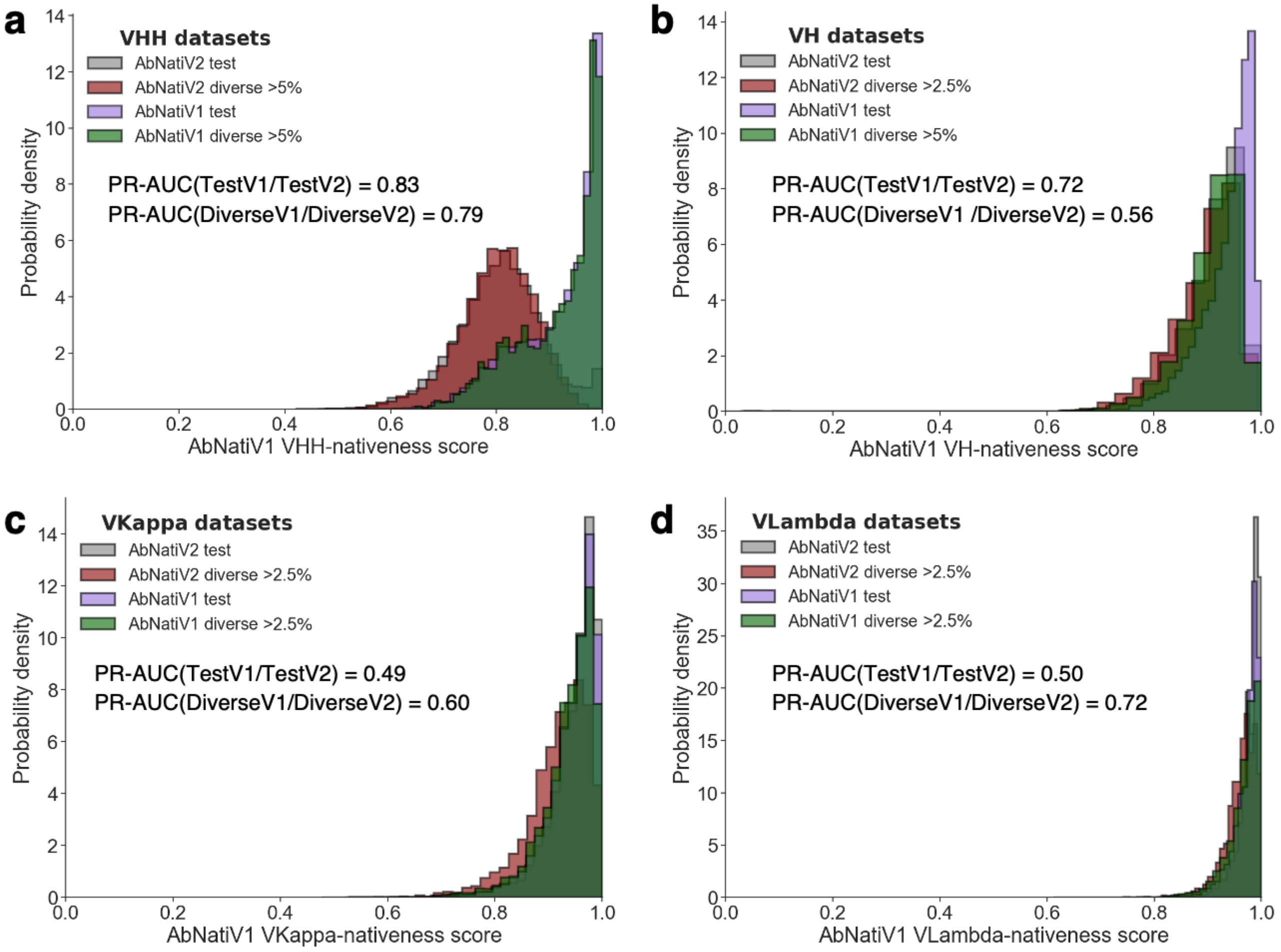


**Supplementary Fig. 8. PR-AUC classification performances of AbNatiV1 between the test and diverse test sets of AbNatiV1 and AbNatiV2**. Nativeness-score distributions and corresponding PR-AUC values used to quantify the ability of AbNatiV1 to distinguish the $V_H$H camelid (**a**), VH human (**b**), VKappa human (**c**), and VLambda (**d**) test and diverse test sets of AbNatiV1 and AbNatiV2 (see legend). Each AbNatiV1 model was trained on ~2million sequences, and hyperparameters are the same between models. However, while human training sequences were highly diverse and came from many different studies, the vast majority of training camelid nanobody sequences only came from three studies, likely explaining the much poorer generalisation of the $V_H$H-AbNatiV1 model compared to the human AbNatiV1models, which score most human sequences ≥ 0.8 as expected.

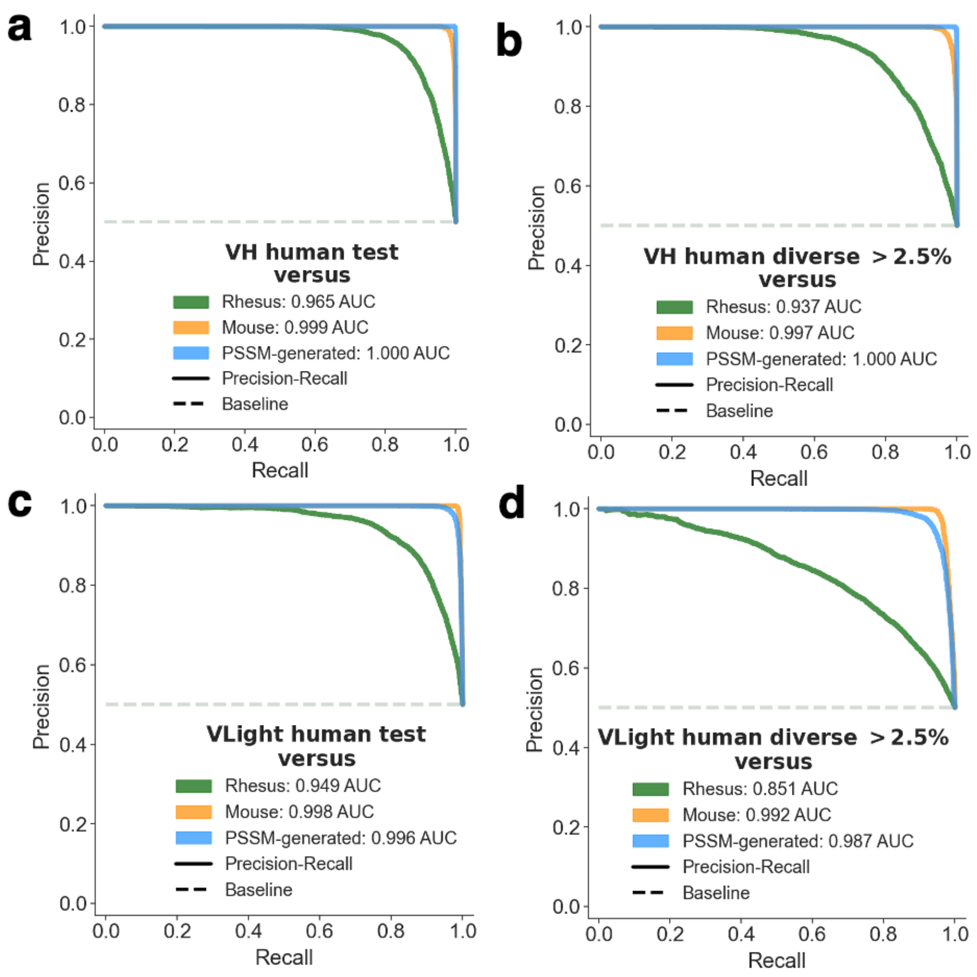


**Supplementary Fig. 9. PR-AUC classification performances for the AbNatiV2 VH and VL models**. Plots of the PR curves used to quantify the ability of AbNatiV2 to distinguish the VH human test (**a**), VH human diverse test (**b**), VL human test (**c**), and VL human diverse test (**d**) sets from the other datasets (see legend, which also reports the AUC values). The baseline (dashed line) corresponds to the performance of a random classifier.

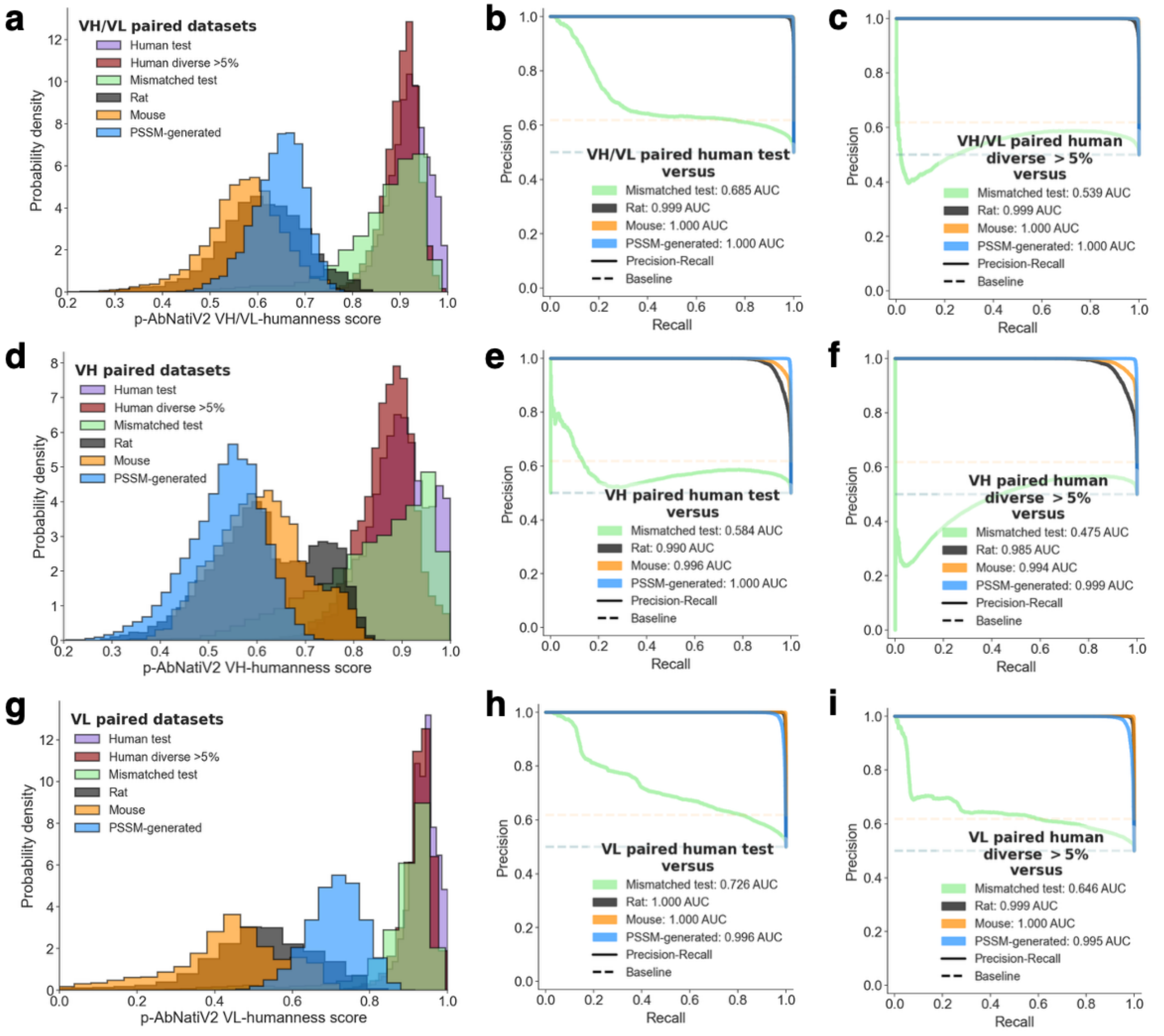


**Supplementary Fig. 10. Performance of the p-AbNatiV2 model on VH/VL paired classification**. (**a, d, g**) The p-AbNatiV2-humanness score distributions of the paired human test (purple), human diverse >2.5% (red), rat (black), PSSM-generated (blue), mouse (orange) VH/VL paired, and mismatched test human sequences generated by mixing heavy and light chains from different B-cell classes and studies (green) (**a**), and heavy (**d**) and light (**g**) chain sequences derived from the corresponding pairs. The PSSM-generated database is made of artificial sequences randomly generated using residue positional frequencies from the PSSM of the human test dataset. The paired human diverse >5% dataset is made of sequences from the test dataset with a sequence identity difference of 5% from their respective closest sequence of the corresponding training set (see Methods). Each dataset contains 10,000 sequences except (**b, c**) Plots of the PR curves computed to represent the ability of p-AbNatiV2 to distinguish the paired human test set (**b**) or paired human diverse >5% (**c**) from the other datasets (see legend, which also reports the area under the curve). (**e, f**) Same PR plots but for the heavy chain sequences derived from the corresponding pairs. (**h, i**) Same PR plots but for the light chain sequences derived from the corresponding pairs. The baseline (dashed line) corresponds to the performance that a random classifier would have.

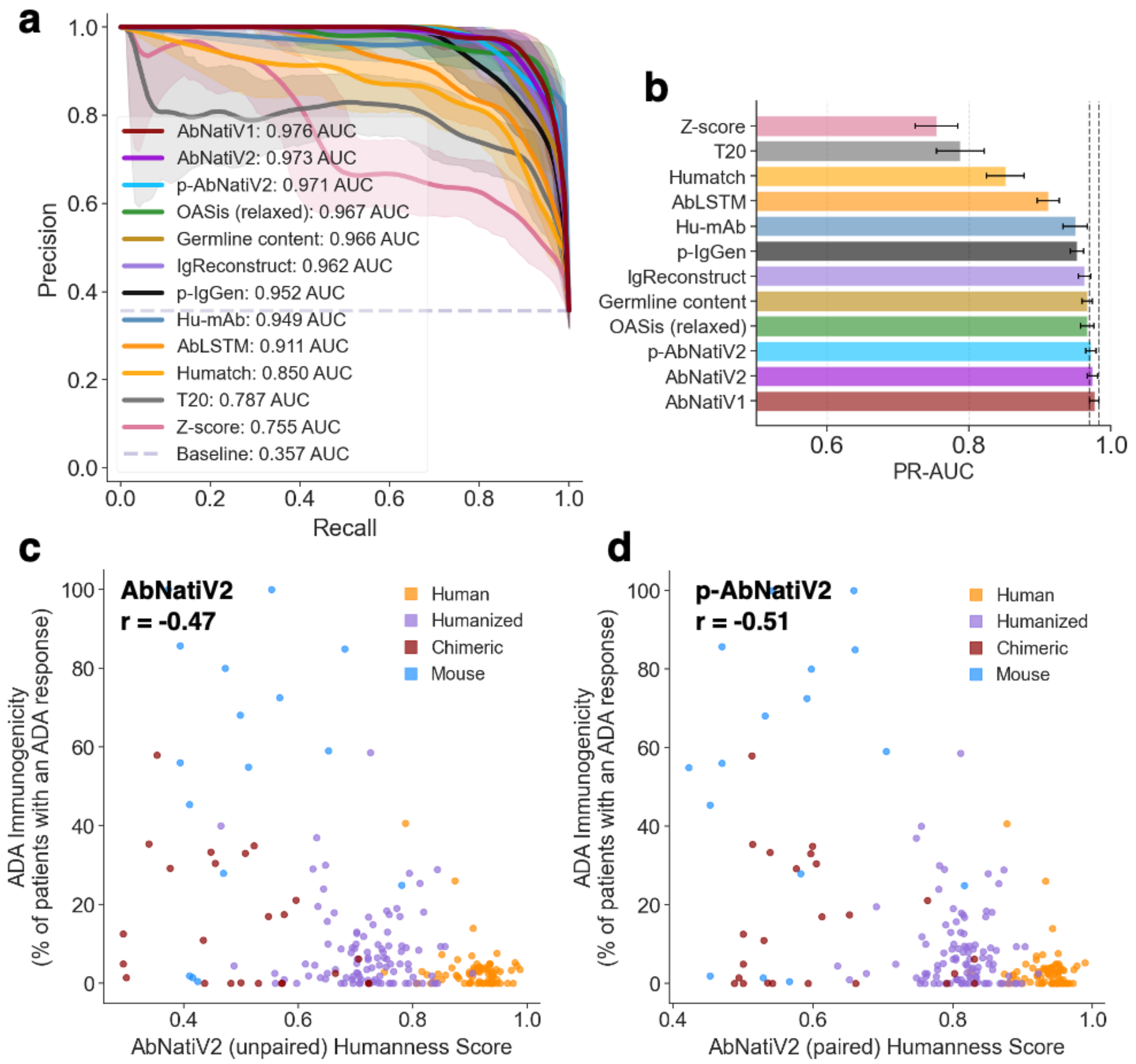


**Supplementary Fig. 11. AbNatiV2 performances on therapeutic antibodies**. (**a**) Plot of the PR curves of the classification of 190 human-derived therapeutics from 342 therapeutics of non-human origin (mouse, chimeric and humanised) carried out with p-AbNatiV2 (in yellow), unpaired AbNatiV2 (in purple), AbNatiV1 (in brown) and nine other computational methods (see legend). PR curves represent the mean of 1,000 bootstrap resamples, and the shaded areas indicate 95% confidence intervals. The legend reports the mean PR-AUC over the bootstrapping. The baseline (dashed line) corresponds to the performance expected from a random classifier. (**b**) Mean PR-AUC values and standard error bars calculated via bootstrapping (1,000 resamples). The top-performing method (AbNatiV1, in red) is annotated with vertical dotted lines at its confidence bounds to facilitate comparison. (**c, d**) Scatter plots of the unpaired AbNatiV2 (**c**), and paired p-AbNatiV2 (**d**) humanness score of 216 antibody therapeutics and their ADA immunogenicity score, expressed as the percentage of patients developing an ADA response in each clinical study. The Pearson correlation (r) is reported on top left corner. Sequences are coloured on the basis of their origin (see legend). For the unpaired AbNatiV2 models, the humanness scores of the heavy and light chains were averaged to yield a single humanness score.

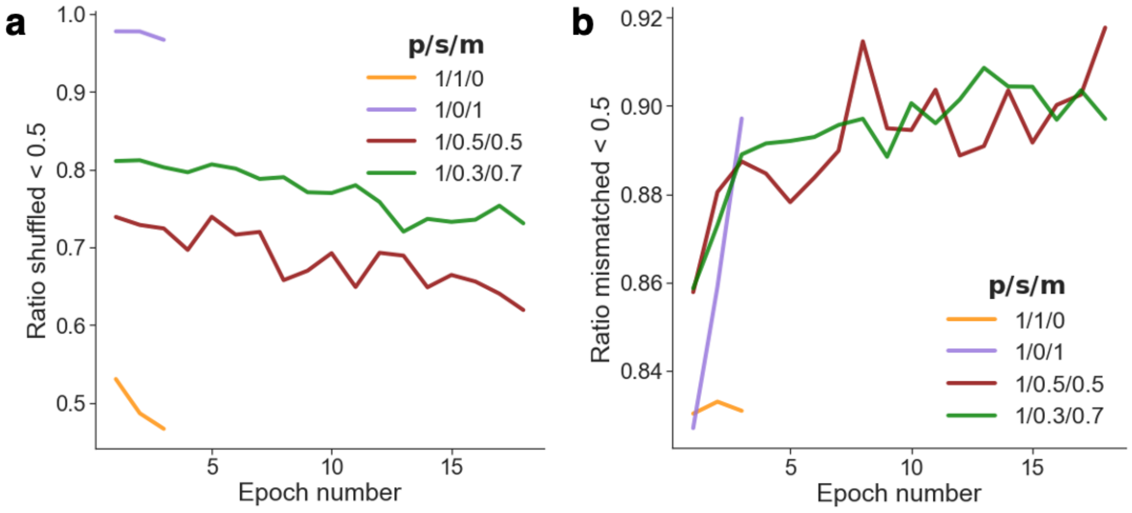


**Supplementary Fig. 12. Impact of paired, shuffled, and mismatched loss weight on p-AbNatiV2 performance under noise-contrastive learning.** The p-AbNatiV2 model was trained with four different paired (p), shuffled (s) and mismatched (m) weight ratios (see legend). (**a**) Ratio of shuffled test sequences with a pairing likelihood below 0.5. (**b**) Ratio of mismatched test sequences with a pairing likelihood below 0.5. The training of underperforming (or very unbalanced) models was terminated after a few epochs.

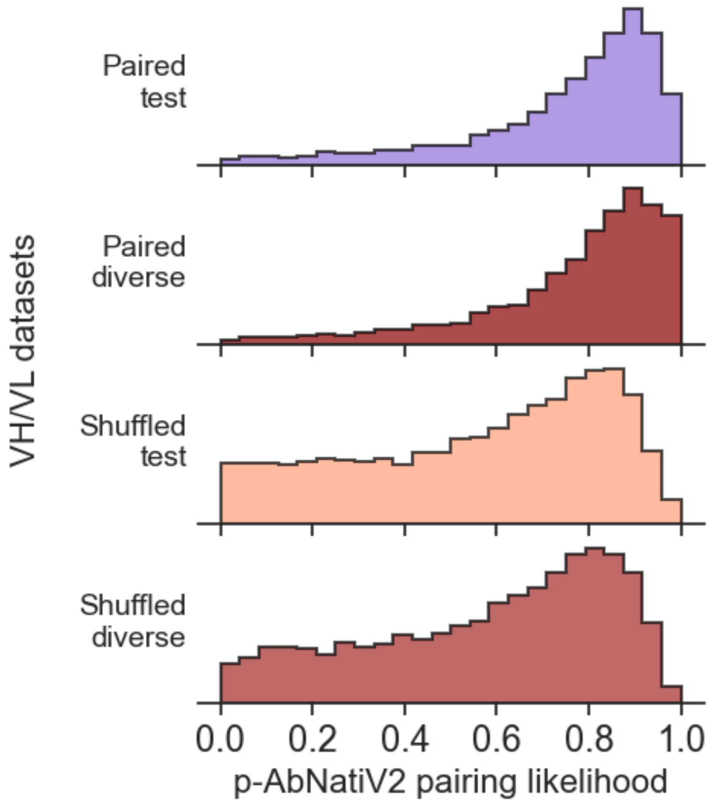


**Supplementary Fig. 13. Distributions of p-AbNatiV2 pairing likelihood scores on the test and diverse test sets**. Distributions of p-AbNAtiV2 pairing scores for 10,000 native human paired test (in purple) and diverse (in red) sequences and shuffled pairs generated by scrambling the light chains across the different native pairs (in beige) and diverse pairs (in salmon). p-AbNatiV2 performs as good on the diverse sets. No overfitting is observed.

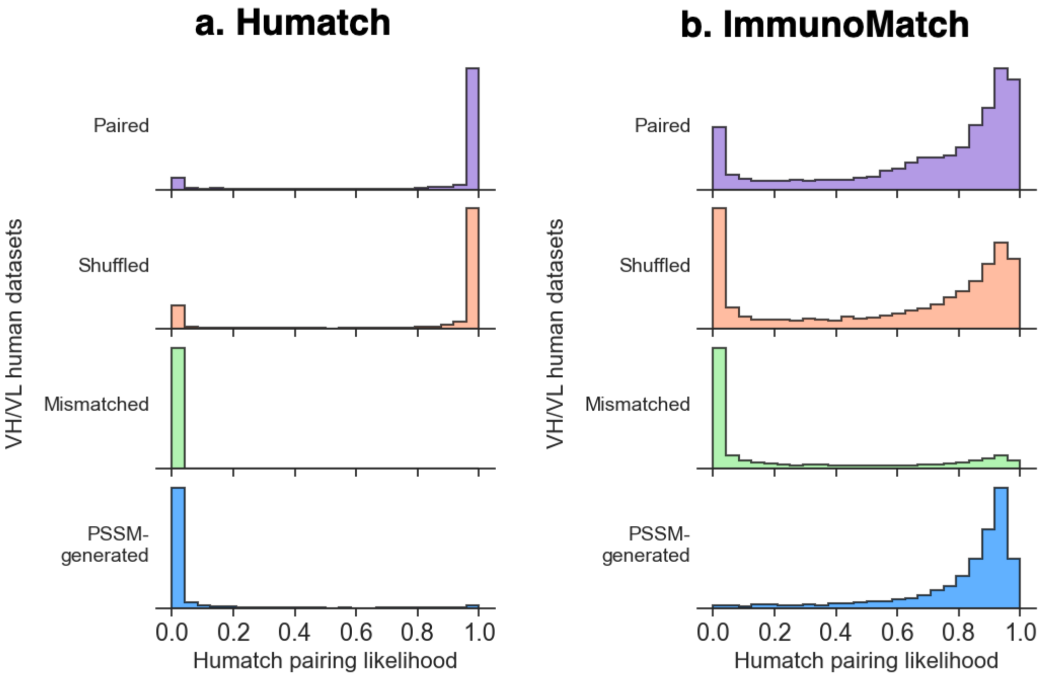


**Supplementary Fig. 14. Distributions of Humatch and ImmunoMatch pairing likelihood scores**. Distributions of Humatch (**a**) and ImmunoMatch (**b**) pairing scores for 10,000 native human paired test sequences (in purple), shuffled pairs generated by scrambling the light chains across the different native pairs within the same somatic maturation bin (in beige), mismatched human sequences generated by mixing heavy and light chains from different B-cell classes and studies (in green, see Methods), and PSSM-generated artificial sequences (in blue). The PSSM-generated database is made of artificial VH and VL sequences randomly generated using residue positional frequencies from the PSSM of the human train dataset.

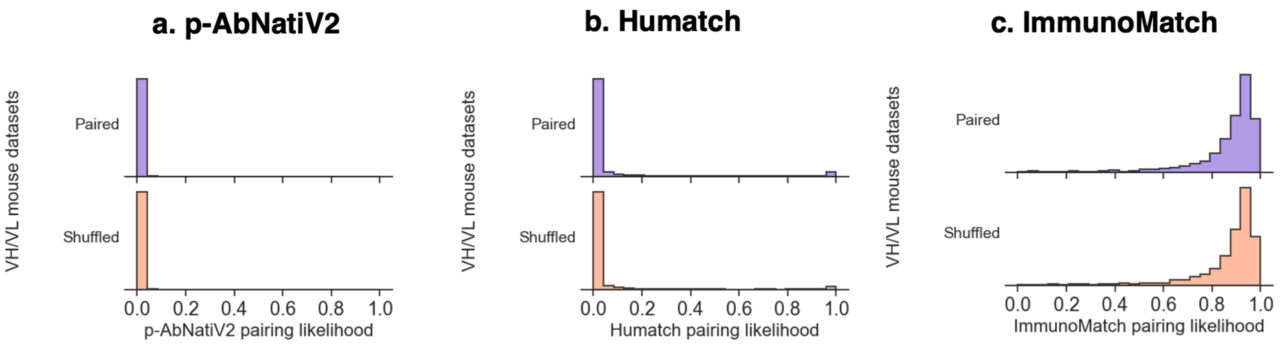


**Supplementary Fig. 15. Pairing likelihood on mouse paired and scrambled Fv sequences**. Distributions of p-AbNatiV2 (**a**), Humatch (**c**) and ImmunoMatch (**d**) pairing scores for 6,971 native mouse paired sequences from the OAS (in purple) and shuffled mouse pairs generated by scrambling the light chains across the different native pairs (in beige).

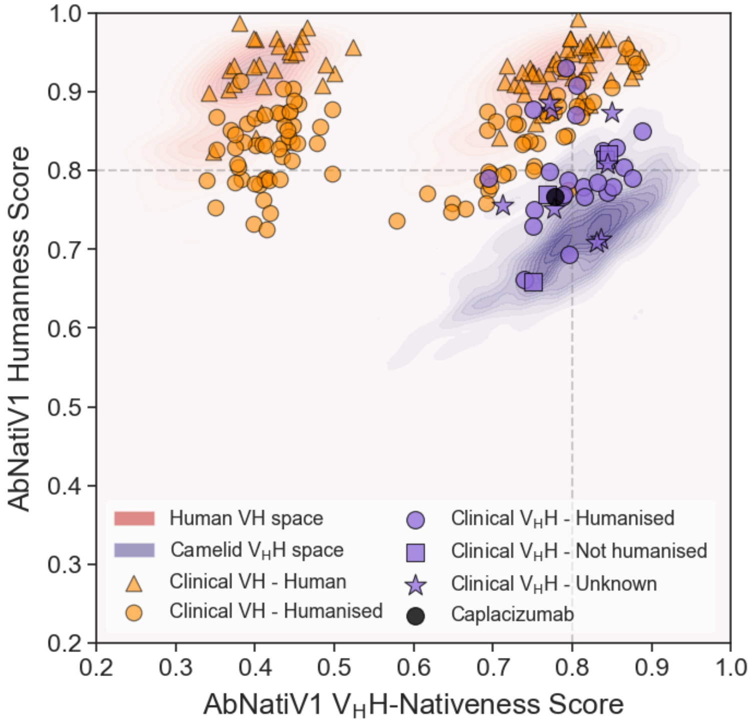


**Supplementary Fig. 16. The nativeness landscape of therapeutic antibodies and nanobodies with AbNatiV1.** AbNatiV1 humanness versus VHH-nativeness scores visualised as kernel densities for 500 sequences from the human VH test dataset (red areas) and camelid VHH test dataset (purple areas). Overlaid are clinical-stage sequences: 170 VH antibodies (orange markers) classified as human (triangles) or humanised (circles) by Prihoda et al., and 36 VHH nanobodies (purple markers) from the Thera-SAbDab dataset classified as humanised (circles), non-humanised (squares), or unknown (stars) based on manual literature curation (see Supplementary Table 4 and Methods). The first FDA-approved nanobody drug, Caplacizumab, is highlighted in black. The corresponding AbNatiV2 landscape is provided in **Figure 3** and the axis ranges are the same for ease of comparison.

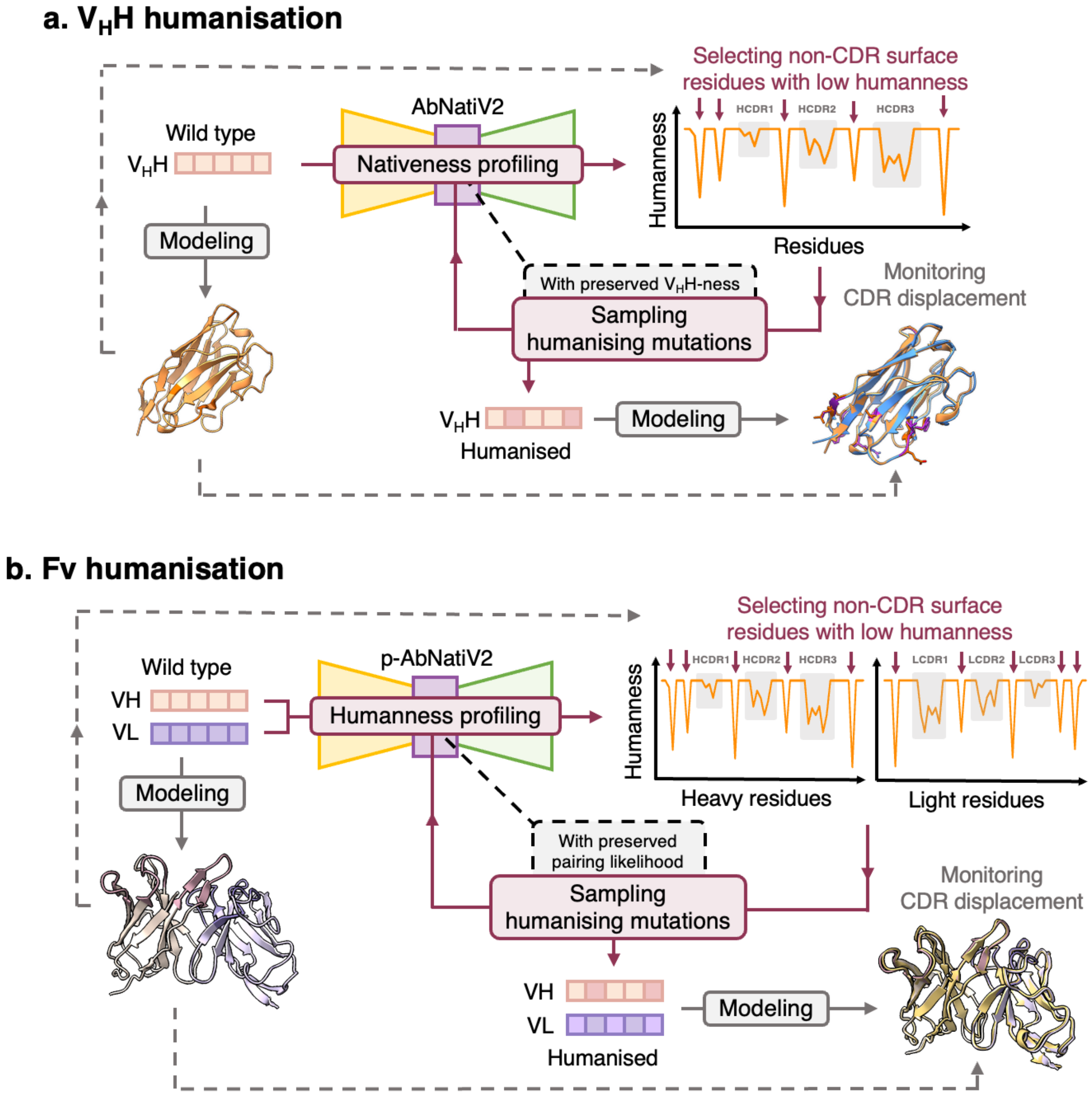


**Supplementary Fig. 17. Illustration of the AbNatiV2 humanisation pipelines**. Humanisation pipelines of $V_H$Hs (**a**) and Fvs (**b**) using AbNatiV2. For both pipelines, the WT sequence is modelled using ImmuneBuilder2 to identify surface-exposed residues. Non-CDR surface residues with low AbNatiV2 humanness are selected for humanisation. Humanising mutations are sampled while preserving VH-VL pairing likelihood (for Fvs) or $V_H$H-nativeness (for $V_H$Hs). Ultimately, the humanised variant is modelled, structurally superimposed to the WT framework model, and CDR displacement is monitored by RMSD to assess the quality of the humanisation.

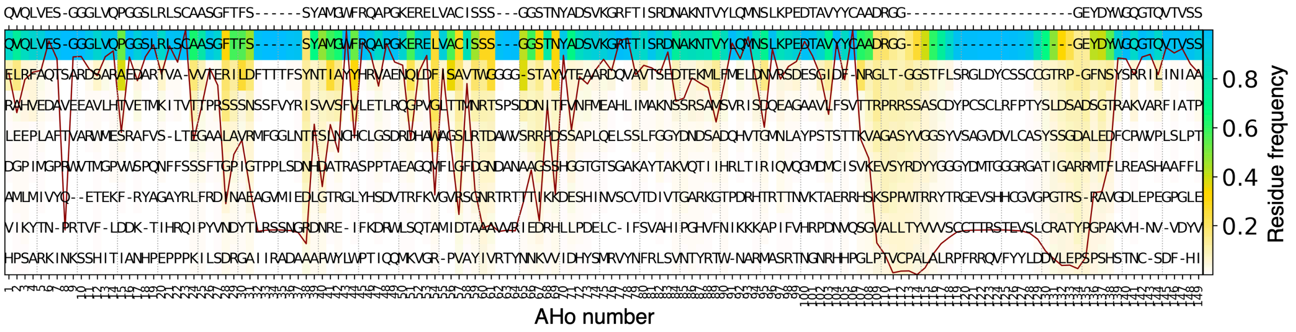


**Supplementary Fig. 18. Position-specific weight matrix (PWM) of a camelid $V_HH$ library amplified on the constant region**. PWM is calculated on the in-house alpaca library sequenced in this study. The sequences were amplified on the leader sequence and the constant region (CH2), then gel-extracted to separate traditional VH (VH+CH1 in gel) from $V_HH$s before deep sequencing. The x-axis reports the AHo numbering scheme used for the alignment, the colour-bar is the amino acid frequency at each position. Each column is sorted from most to least frequent residue at that position, and only the top eight residues are shown. Residues are coloured based on their frequency at each position (see colour-bar). The consensus sequence, which consists in the most frequent residue at each position, is shown on the top. The continuous red line is the conservation index of each position (high means position highly conserved, low poorly conserved).

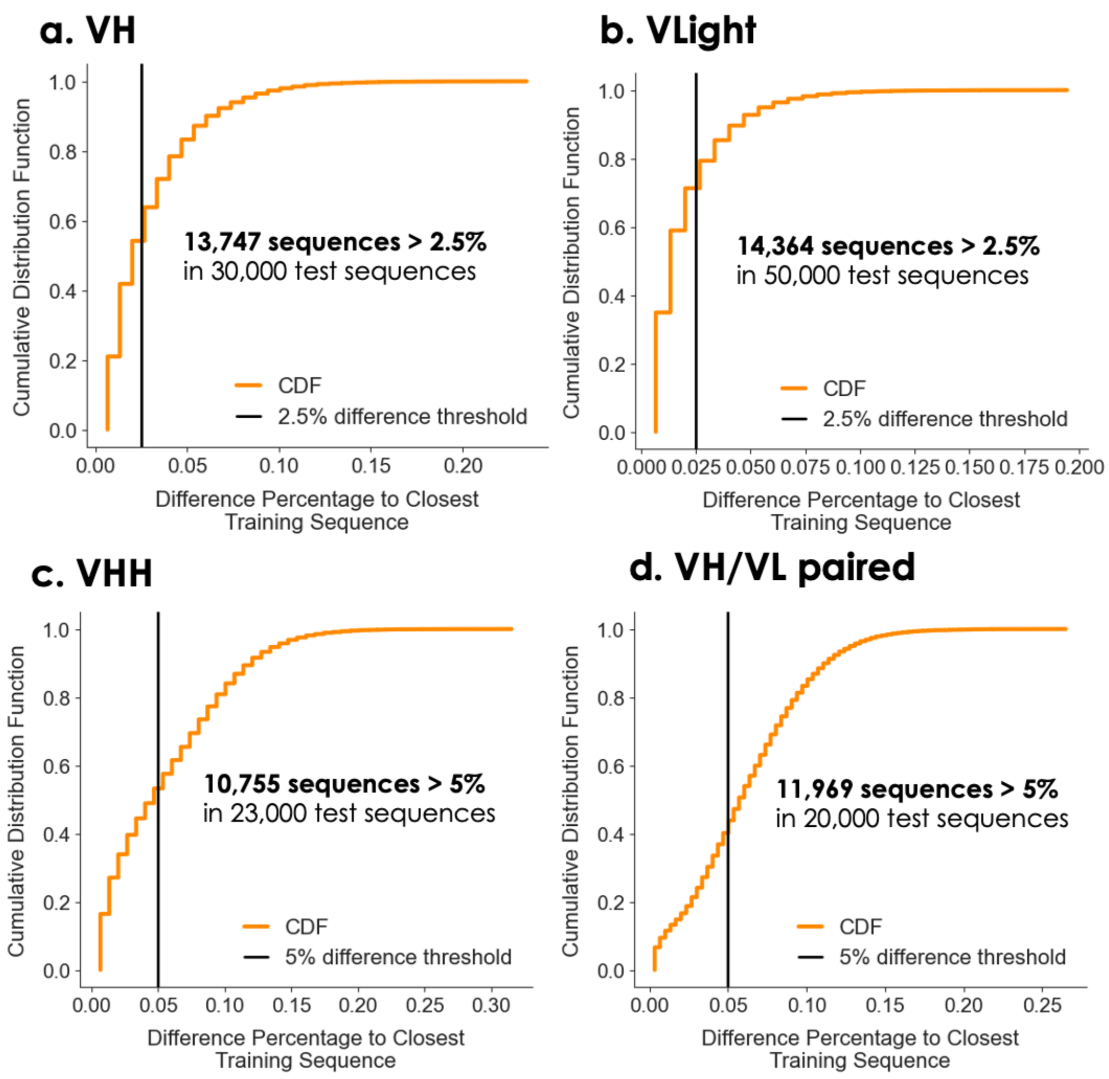


**Supplementary Fig. 19. Minimum percent difference from training sequences.** Each panel reports the cumulative distribution functions (CDFs) for the VH Human Test (**a**), VLight Human Test (**b**), $V_HH$ Camelid Test (**c**), and VH/VL paired Human Test (**d**). The x-axis reports the percent difference to the closest sequence in the respective training dataset. The black line is the minimum difference percentage threshold used to extract, for each AbNatiV model, the corresponding Diverse test set. For each of these sequences, the calculation is done against ~20 million sequences in the training set, making it computationally very expensive. For the VLight sequences, the distribution was computed on the whole test dataset of 50,000 sequences. For VH, $V_HH$ the calculation was performed only on 30,000 sequences and 23,000 sequences for VH/VL paired VH+VL as they already led to more than 10,000 diverse sequences. For the paired sequences, the heavy and light chains were concatenated prior to distance calculations.

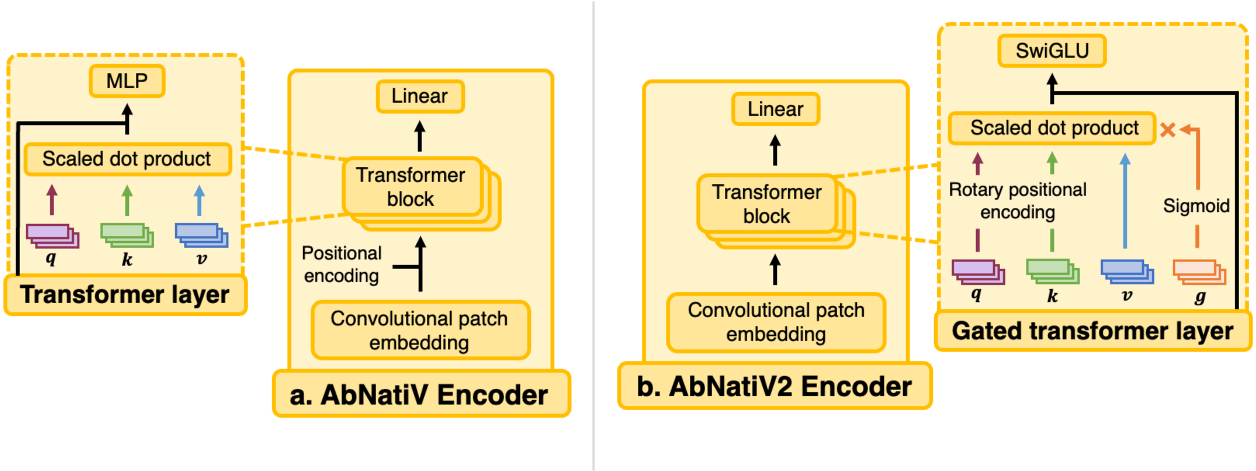


**Supplementary Fig. 20. Architecture comparison between AbNatiV and AbNatiV2**. Schematic representation of the encoder architectures of AbNatiV (**a**) and AbNatiV2 (**b**). In AbNatiV2, the sinusoidal positional encoding is replaced with a rotary positional encoding, applied to the queries $q$ (in burgundy) and keys $k$ (in green) within the transformer layers to enhance positional information. The scaled dot product attention mechanism, which integrates the queries $q$ , keys $k$, and values $v$ (in blue), is further modulated by an additional gate $g$ (in orange) in each layer. This gate dynamically scales the output of the dot product using a sigmoid activation function. Furthermore, the Multi-Layer Perceptron (MLP) in the residual connection is replaced by a SwiGLU activation mechanism, which improves efficiency and representation capacity. The same updates are applied to the decoder

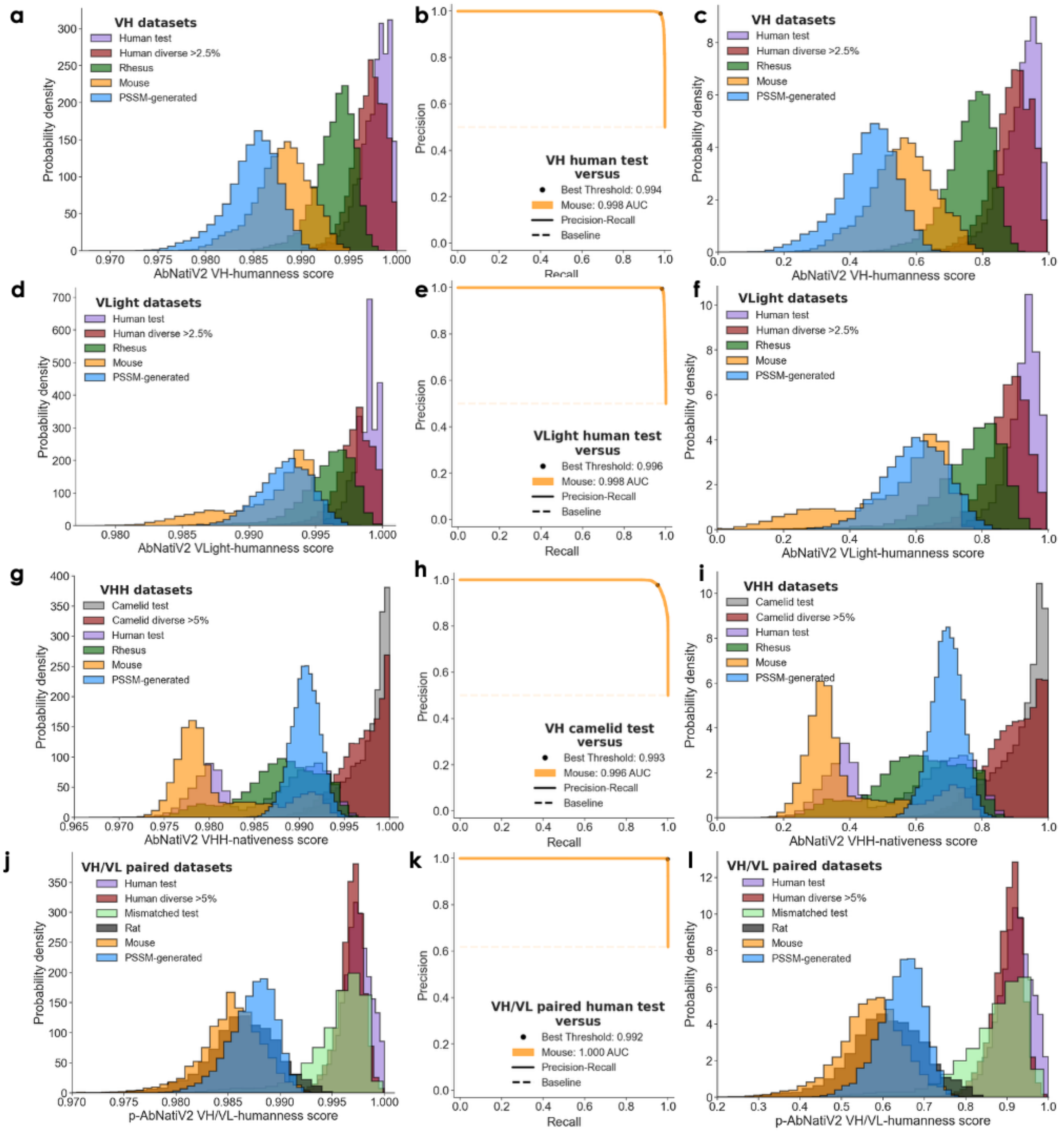


**Supplementary Fig. 21. Linear rescaling of the AbNatiV2 nativeness score**. (**a, d, g, j**) The AbNatiV2 nativeness score distributions of various datasets (see legend) after the $\exp(-MSE)$ projection but before the linear rescaling for models trained on human VH (**a**), VLight (**d**), camelid $V_HH$ (**g**), VH/VL paired (**j**) sequences. The VLight set combines the VKappa and VLambda test sets together. (**b, e, h, k**) The optimal thresholds $T_R$ (black marker) extracted as the point closest to (1,1) in the PR curves when separating the Human Test sequences from the Mouse sequences for the models trained on human VH (**b**), VLight (**e**), and VH/VL paired (**k**) sequences, and when separating Camelid Test sequences from the Mouse sequences for the model trained on camelid $V_HH$ sequences (**h**). (**c, f, i, l**) The AbNatiV2 nativeness score distributions (respectively from **a, d, g,** and **j**) after linear rescaling $((0.8 - 1) * (X - 1)/(T_R - 1) + 1)$ for each model following their respective $T_R$ (respectively from **c, f, i,** and **l**). The same procedure was applied separately to the heavy and light chain assessments of the paired model. This rescaling ensures that a nativeness score of 0.8 is the threshold that best separate native from non-native sequences across all models.
